# Control of Synaptic Communication through Molecularly Engineered Bioluminescence Light Emission and Sensing

**DOI:** 10.1101/2025.10.27.684942

**Authors:** Ashley N. Slaviero, Mansi Prakash, Rachel Schumaker, Emmanuel L. Crespo, Alexander D. Silvagnoli, Maya O. Tree, Gerard G. Lambert, Christopher I. Moore, Nathan C. Shaner, Diane Lipscombe, Ute Hochgeschwender

## Abstract

Synapses are sites of intercellular communication between neurons and from neurons to target organs, and of signal integration that underly physiological and behavioral responses. We have developed a modular platform, Interluminescence (Int), for experimental control of synaptic transmission: bioluminescent light, generated by a luciferase oxidizing a luciferin, from a presynaptic neuron is used to activate transsynaptic optogenetic ion channels in the postsynaptic neuron. Two strategies can modulate postsynaptic neurons in the presence of luciferin. In the ‘Act-Int’ approach, a luciferase is genetically expressed in synaptic vesicles and released during depolarization-induced presynaptic vesicle fusion and exocytosis. In the ‘Persist-Int’ approach, a luciferase is tethered to the presynaptic membrane where it can support sustained transsynaptic signaling. Both strategies can activate postsynaptic neurons with comparable efficacy under the conditions tested. By design, the modularity of the platform permits the use of luciferases and opsins ranging in brightness and light sensitivity, with the luciferase targeted to different subcellular regions of the presynaptic neuron, and the postsynaptic opsin being excitatory or inhibitory. Our results demonstrate the utility and versatility of Interluminescence to mediate synapse-specific transmission that is either activity-dependent or activity-independent.

## Introduction

Genetic tools have revolutionized neuroscience research by making it possible to dissect neural networks with temporal and spatial precision (e.g., opto-, chemo- and sonogenetics^1–4^). Through genetically targeted cell-specific expression of actuators, it is now possible to use light, chemicals, or ultrasound to activate specific neuronal circuits and measure the resulting change in activity, or behavior. There are nonetheless gaps in the ability to manipulate and override neural communication, specifically the research community lacks tools that permit direct experimental control of the efficacy and form of synaptic transmission between specific partners.

We and others have developed several methods using bioluminescence as a local source of light for optogenetic control of neural circuits^5–12^. Several recent advances in molecular engineering and the development of brighter luciferases have expanded the utility of bioluminescence for optical control in neurons both *in vitro* and *in vivo*^13–15^. Here, we capitalize on the fact that both light-generating (luciferase) and light-sensing (opsin) components are genetically encoded and thus can be expressed in separate genetically determined locations. Expressing the luciferase in a presynaptic neuron and the opsin in a postsynaptic neuron creates an “optical synapse”; information is transmitted by bioluminescence-dependent activation of opsins across defined synaptic partners, enabled by the presence of luciferin. This transsynaptic form of bioluminescence signaling - Interluminescence (Int) - allows for non-invasive, temporal control of signaling across synapses of neural circuits deep within the brain. Furthermore, Interluminescence could be used to create *de novo* transsynaptic signaling, effectively re-wiring neural circuits for experimental inquiry or to compensate for hypo or hyper-active synapses.

Recently we showed that luciferases could be packaged in presynaptic vesicles, released along with neurotransmitters and – in the presence of luciferin – activate opsins expressed postsynaptically^16^. We expand on activity-dependent Interluminescence (Act-Int) by developing bioluminescent transsynaptic signaling that is persistent and independent of presynaptic activity (Persist-Int). **Figure 1** highlights key features of the Interluminescence strategies in comparison to a classical neurotransmitter mediated synapse.

**Fig. 1.**
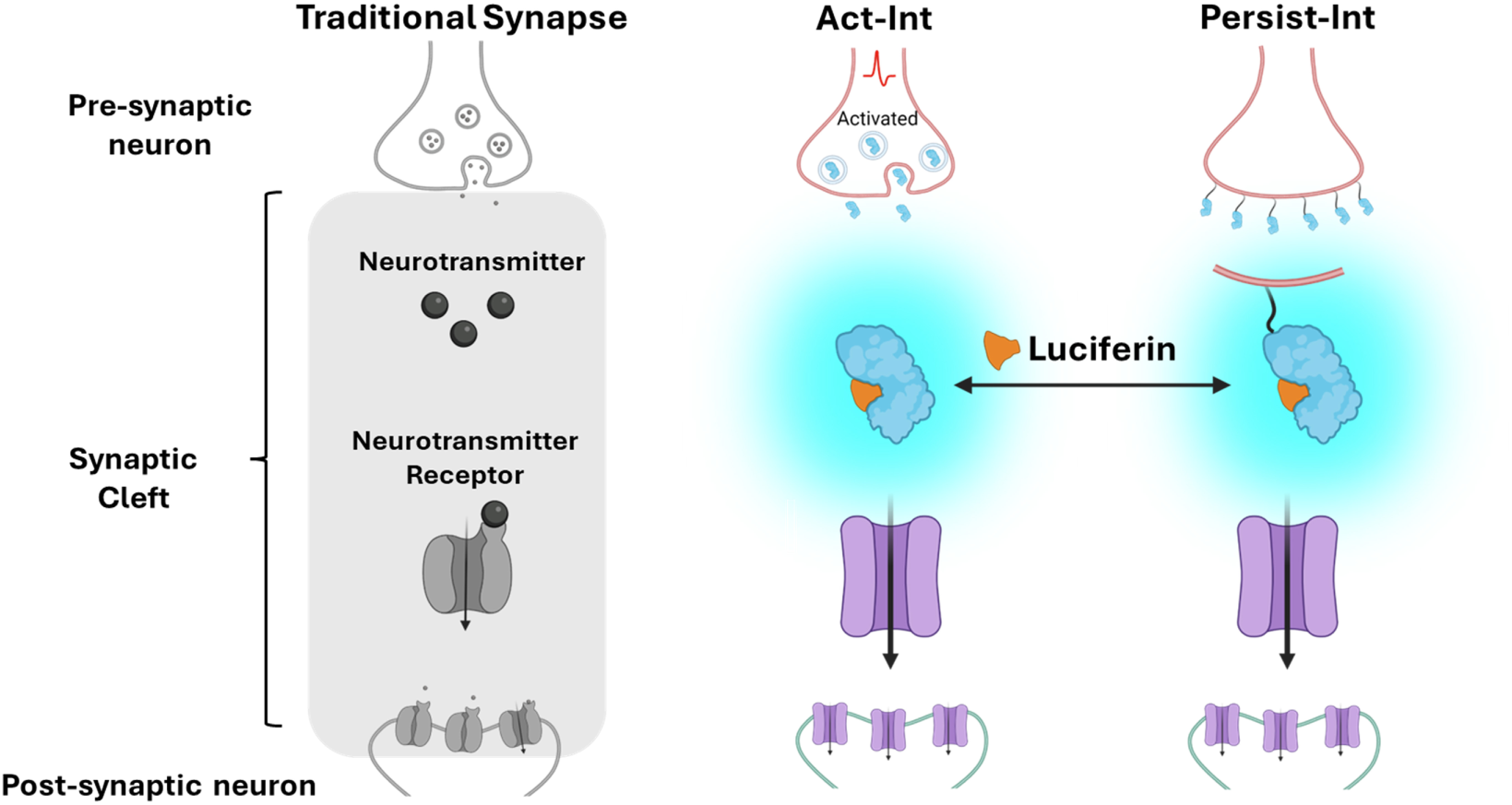
Transient activity-dependent and persistent activity-independent forms of Interluminescence. Neurotransmitter-mediated synaptic transmission is compared to activity-dependent (Act-Int) and activity-independent (Persist-Int) forms of Interluminescence. By comparison to neurotransmitter-mediated synaptic transmission, Interluminescence depends on light emitted from a luciferin-luciferase chemical reaction to travel across the synaptic cleft and activate an opsin expressed in the postsynaptic cell. In Act-Int, the luciferase is in synaptic vesicles and released (along with neurotransmitter) during synaptic vesicle exocytosis. In Persist-Int, the luciferase is in the synaptic cleft anchored to the extracellular face of the presynaptic cell. Persist-Int is independent of presynaptic vesicle release and is activated whenever the luciferin is present, whereas Act-Int depends on presynaptic vesicle release and the presence of the luciferin. Compared to neurotransmission, which is mediated by neurotransmitter diffusion across the synaptic cleft and activation of postsynaptic neurotransmitter receptors, Persist-Int and Act-Int are independent of neurotransmitters, but both require luciferin, and the resulting postsynaptic response depends on the nature of the opsin expressed.

Here we demonstrate efficient synaptic transmission with the Persist-Int approach, observing activation and inhibition of postsynaptic neurons in multiarray electrode recordings in culture. We compare postsynaptic responses mediated by Act-Int and Persist-Int to neurotransmitter (NT)-mediated postsynaptic events. We further show that for both approaches light emitters and light sensors ranging in brightness and light sensitivity can be combined to tune the gain and polarity of synaptic transmission. Lastly, we demonstrate the synapse specificity of both Int approaches.

## Results

### Activity-independent Interluminescence: Persist-Int

We showed previously that Interluminescence could be used for activity dependent transsynaptic signaling in cultured neurons by packaging luciferases in presynaptic vesicles^16^. This form of Interluminescence requires presynaptic depolarization and vesicle exocytosis (Act-Int; **Fig. 1**). Here we develop an activity-independent form of Interluminescence mediated by a luciferase-luciferin reaction within the synaptic cleft (Persist-Int; **Fig. 1**).

We first tested Persist-Int in cultured neurons. To target the luciferase to the extracellular face of the presynaptic membrane, we used a fusion protein of mNeonGreen and the NanoLuc variant eKL9h (GeNL_eKL9h, a bright molecularly evolved FRET, Förster resonance energy transfer, construct^15^) connected to ICAM-Nrxn3b^17^ via the 15 amino acid linker used in luminopsins^5^. Nrxn3b (a pre-synaptic adhesion protein neurexin variant) tethered GeNL_eKL9h to the extracellular presynaptic membrane, and the extracellular domain of ICAM-1 (a cell surface expressed glycoprotein adhesion receptor) was included to extend the light emitter further into the synaptic cleft^17,18^. The complete presynaptic luciferase construct, GeNL_eKL9h-ICAM-Nrxn3b-P2A-dTom, contained the red fluorescent protein dTomato for visual identification of the fusion protein in neurons (**Fig. 2A**). The inhibitory anion channel hGtACR2 and the excitatory step function opsin ChR2(C128S), were selected as the postsynaptic opsins to achieve optimal activation by blue light^19,20^. Both opsin constructs contained the yellow fluorescent protein EYFP for visual confirmation of expression in neurons.

**Fig. 2.**
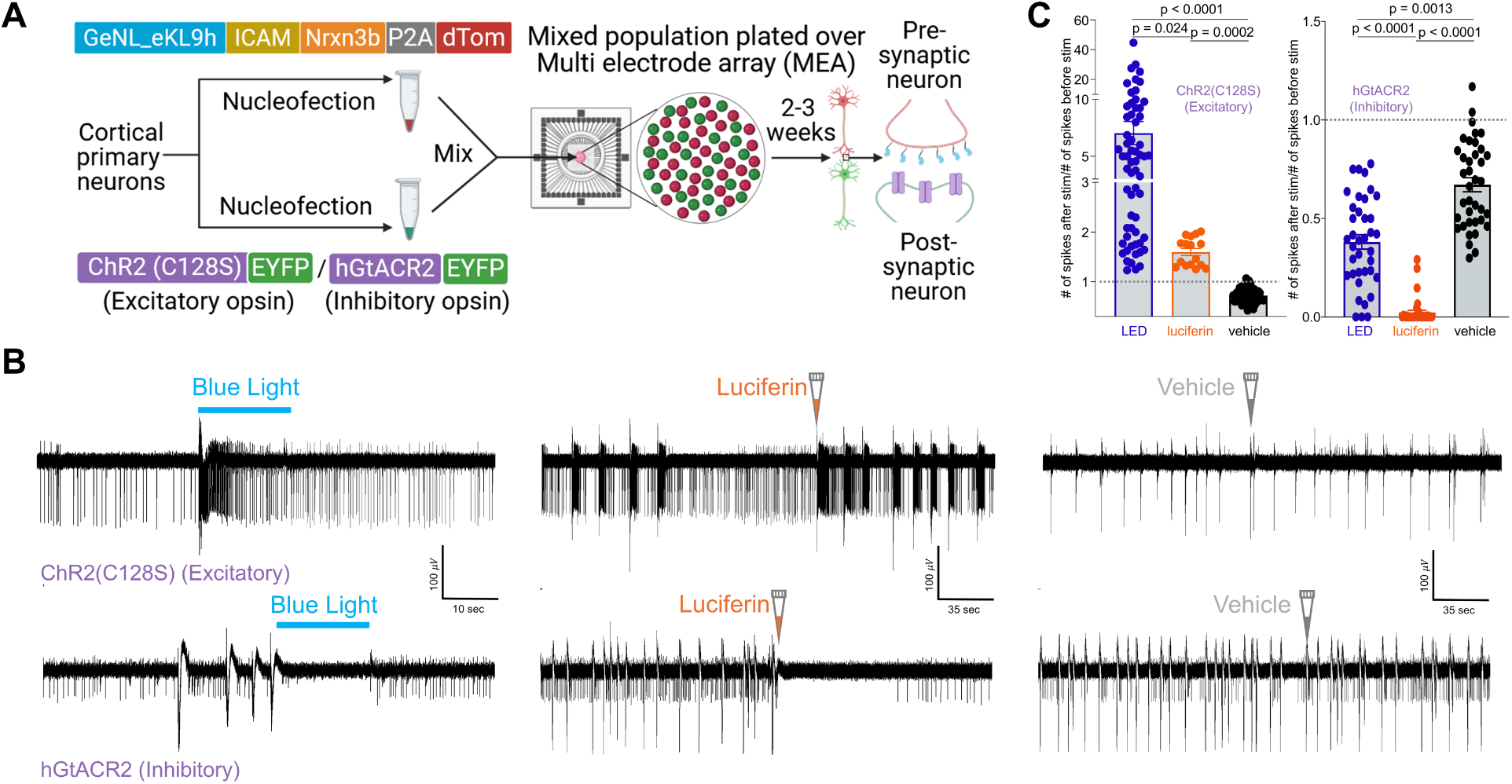
Modulation of postsynaptic neural activity by Persist-Int in vitro. **A** Schematic of experimental design and constructs used in cortical neuron nucleofections with either one of the opsins, ChR2(C128S)-EYFP (excitatory) and hGtACR2-EYFP (inhibitory), or GeNL_eKL9h-ICAM-Nrxn3b-P2A-dTomato. Neurons were mixed (red and green spheres), plated on MEAs, and maintained in culture for 2-3 weeks until recording. **B** Representative traces from individual electrodes of MEAs with postsynaptic neurons expressing the excitatory opsin ChR2(C128S) (upper panels) and the inhibitory opsin hGtACR2 (lower panels) showing response to blue light, luciferin (hCTZ; 10 μM)-induced bioluminescence and vehicle. **C** Scatter plots combining data from individual electrodes across multiple experiments showing signal-to-noise ratios derived from the spike frequency 5 sec after stimulation divided by the spike frequency 5 sec before stimulation. Left panel is excitatory opsin expressing postsynaptic neurons and right panel is inhibitory opsin expressing postsynaptic neurons as depicted for individual traces in B. ChR2(C128S): N= 62 (LED), 16 (Luciferin), 53 (Vehicle); hGtACR2: N = 38 (LED), 38 (Luciferin), 38 (Vehicle); Kruskal-Wallis; Excitatory: LED vs Luciferin p = 0.024, LED vs Veh p < 0.0001, Luciferin vs Veh p; Inhibitory: LED vs Luciferin p < 0.0001, LED vs Veh p = 0.0013, Luciferin vs Veh p < 0.0001.

We tested the ability of luciferase, tethered to the extracellular surface of the presynaptic membrane, to generate sufficient local excitation upon exposure to the luciferin to activate postsynaptic opsins in co-cultures of cortical neurons (**Fig. 2A**). We used nuclear transfection to express the luciferase GeNL_eKL9h-ICAM-Nrxn3b-P2A-dTom construct into one batch of cortical neurons and to express either hGtACR2 or ChR2(C128S) into a separate batch of cortical neurons. We then generated co-cultures of cortical neurons expressing GeNL_eKL9h-ICAM-Nrxn3b-P2A-dTom and either the inhibitory hGtACR2 opsin or the excitatory ChR2(C128S) opsin. Co-cultures were plated on multi electrode arrays (MEAs) to record neuronal activity (**Fig. 2**).

To confirm opsin responsiveness, we exposed MEAs directly to blue LED light and compared these responses to luciferin-induced activity in co-cultures expressing ChR2(C128S) (**Fig. 2B** upper panel) or hGtACR2 (**Fig. 2B**, lower panel). We used the luciferin hCoelenterazine (hCTZ) to initiate the bioluminescence response or vehicle alone as a control. Postsynaptic responses were measured in cultures in response to both direct opsin activation via LED light or application of hCTZ (Persist-Int). The activity of ChR2(C128S) expressing co-cultures was increased, and the activity of hGtACR2 expressing cultures decreased relative to baseline (**Fig. 2B**). The relative change in spike frequency for all recordings under the different experimental conditions is summarized in **Fig. 2C**. In this set of MEA experiments, vehicle application, and thus simply the application of a solution, non-specifically dampened firing activity in neurons expressing both excitatory and inhibitory opsins, though to a lesser extent than by bioluminescence-mediated hGtACR2 activation. In subsequent whole cell recordings, we did not see changes in membrane potential following vehicle/solvent application (see below). The MEA experiments establish that luciferases tethered to the extracellular presynaptic membrane can, in the presence of a luciferin, drive activation of postsynaptic opsins, thereby enhancing or reducing neural activity, depending on the type of target opsin expressed. While quantitatively different (N = 62 electrodes (LED), 16 electrodes (luciferin), 53 electrodes (vehicle); Kruskal-Wallis test: excitatory opsin: vehicle vs LED p < 0.0001, vehicle vs luciferin p = 0.0002, luciferin vs LED p = 0.024; inhibitory opsin: vehicle vs LED p = 0.0013, vehicle vs luciferin p < 0.0001, luciferin vs LED p < 0.0001), the direction of the change in postsynaptic activity following luciferin application as compared to direct LED stimulation was the same between methods (**Figs. 2B, 2C**).

To gain insight into the potential underlying mechanisms of the observed Persist-Int effect on postsynaptic responses, we estimated photon flux from individual emitters^21^ assuming isotropic emission and compared these values to literature irradiance benchmarks for commonly used opsins^19,20,22–27^ (**Supplementary Tables 1 and 2**). Across all emitter–opsin pairs, isotropic single-molecule emission falls short of reported activation benchmarks by several orders of magnitude, even at nanometer-scale distances. These estimates therefore represent a lower-bound radiative comparison and indicate that freely propagating photons from individual emitters are unlikely to account for the observed postsynaptic responses. Instead, they define a regime in which productive signaling must depend on local factors not captured by the isotropic model, including nanoscale proximity, molecular organization, and time integration by the opsin.

To examine whether local coupling could support efficient signaling at synaptic distances, we next constructed a simplified bounding-case model of short-range transfer under favorable geometric assumptions (**Table 1**). In this model, donor–opsin separation was assumed to be on the order of ∼5 nm, yielding high transfer efficiency and maximal coupling. Estimated donor requirements were calculated for the major Persist-Int donor-opsin pairings tested in this manuscript, restricting donors to ICAM-Nrxn3b fusions. FRET efficiency was computed as E = 1 / [1 + (r / R₀)^6^] with R₀ = 5 nm and r = 5 nm, giving E = 0.50. The transferred excitation rate per donor was then taken as E × Vmax. To estimate the number of donor molecules required to reach the opsin benchmark in this FRET-coupled scenario, we inferred an effective capture cross-section from the same operational benchmark framework used in **Supplementary Tables 1 and 2**, which yields N = 1 / (E × Vmax × τoff). Values below 1 are reported as <1 and indicate that a single donor-opsin pair would be theoretically sufficient under this simplified model. Radiative photon transfer was excluded from this calculation. Under these conditions, the number of donor molecules required to achieve opsin activation is small, in some cases approaching unity. These values should be interpreted as optimistic lower bounds that are highly sensitive to distance, orientation, and spectral overlap, and are intended to define a feasible operating regime rather than to predict actual donor requirements in the synapse.

**Table 1.** Estimated donor requirements for FRET-coupled Persist-Int activation

| Persist-Int donor | Donor output at Vmax<br>(ph s<sup>-1</sup> molecule<sup>-1</sup>) | Postsynaptic opsin | τ<sub>off</sub> used (s) | FRET efficiency E | Transferred excitations per donor (s<sup>-1</sup>) | Estimated donor molecules required N* |
| --- | --- | --- | --- | --- | --- | --- |
| NanoLuc-ICAM-Nrxn3b | 8.6 | ChR2(C128S) | 106 | 0.50 | 4.30 | <1 |
| GeNL_eKL9h-ICAM-Nrxn3b | 21.5 | ChR2(C128S) | 106 | 0.50 | 10.75 | <1 |
| GeNL_eKL9h-ICAM-Nrxn3b | 21.5 | hGtACR2 | 0.01 | 0.50 | 10.75 | 9.30 |
| GeNL_eKL9h-ICAM-Nrxn3b | 21.5 | ChRmine | 0.3 | 0.50 | 10.75 | <1 |
| GeNL_eKL9h-ICAM-Nrxn3b | 21.5 | CheRiff | 0.016 | 0.50 | 10.75 | 5.81 |
| GeNL_SS-ICAM-Nrxn3b | 86.0 | ChR2(C128S) | 106 | 0.50 | 43.00 | <1 |
| GeNL_SS-ICAM-Nrxn3b | 86.0 | ChrimsonR | 0.0158 | 0.50 | 43.00 | 1.47 |

\* N refers to concurrently active luciferase donor molecules. Because each catalytic turnover consumes one luciferin substrate molecule, substrate consumption scales directly with donor number and turnover rate, but the table reports donor molecules rather than substrate molecules.

### Interluminescence-dependent postsynaptic action potentials

Having demonstrated Interluminescence by targeting luciferases to the synaptic cleft anchored to the presynaptic membrane (Persist-Int) or to synaptic vesicles (Act-Int), we next compared the properties of postsynaptic action potentials (APs) triggered by Interluminescence and neurotransmitters. We used whole cell recording to monitor postsynaptic membrane potentials in co-cultures of presynaptic cortical (luciferase-expressing) and postsynaptic striatal (opsin-expressing) neurons. The basic experimental design is shown in **Figs. 3A, 3B**. Cortical neurons were nucleofected with the presynaptic constructs for Act-Int (pcDNA-CAG-POMC sbGLuc-P2A-Chrimson-dTom) or Persist-Int (pcDNA-CAG-GeNL_eKL9h-linker-ICAM-Nrxn3b-P2A-dTom), and striatal neurons were nucleofected with the construct for the excitatory opsin ChR2(C128S) (pcDNA CAG ChR2(C128S) EYFP), unless indicated otherwise. Postsynaptic neurons were identified in co-cultures for recording by the presence of EYFP (**Fig. 3A**).

**Fig. 3.**
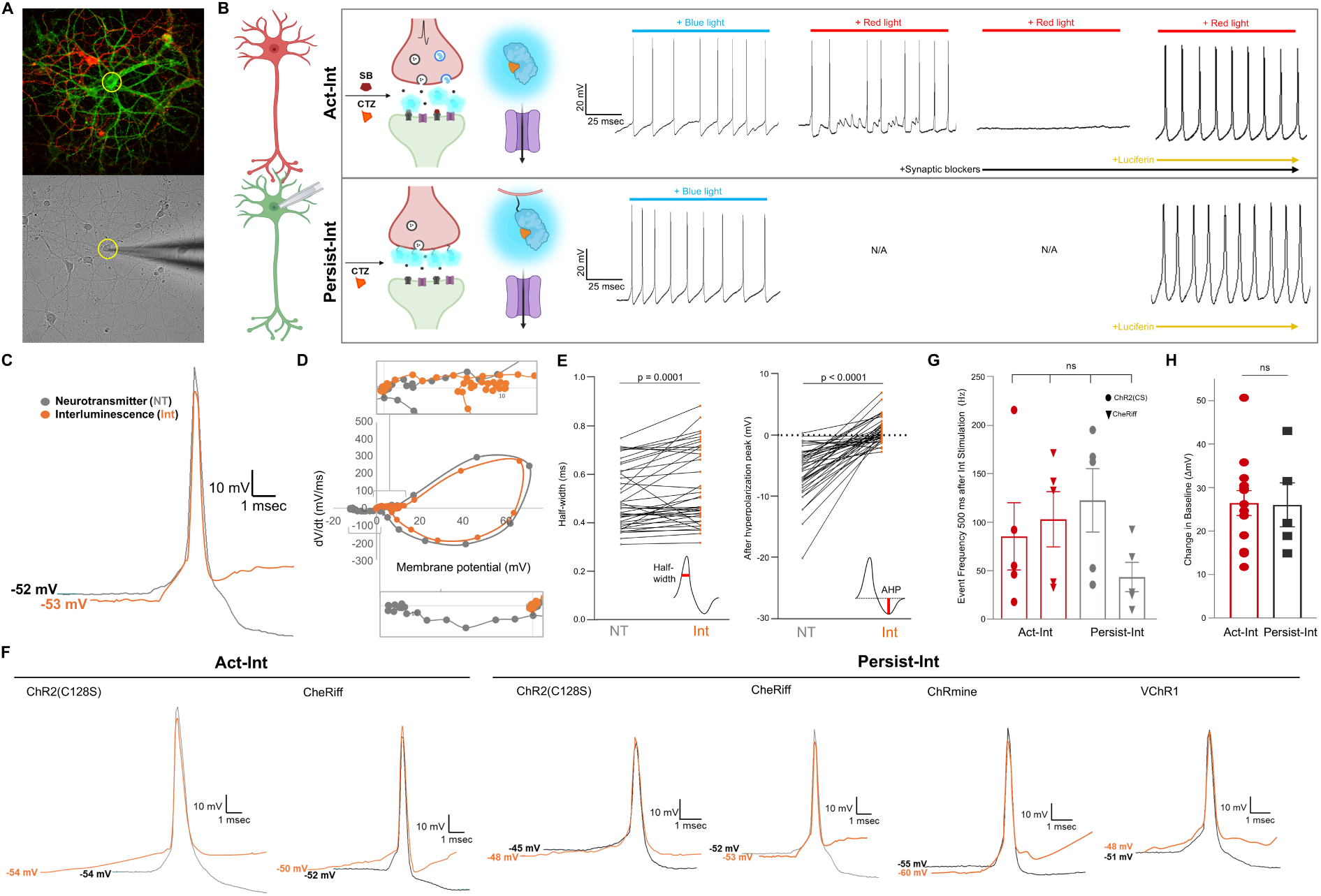
Interluminescence-dependent postsynaptic action potentials. **A** Top: Fluorescent images of Interluminescent co-cultures (red presynaptic, green postsynaptic). Bottom: Brightfield image of patch pipet on postsynaptic neuron. **B** Schematics of Interluminescent synapses at increasing magnification. Patch traces show workflow for postsynaptic Interluminescent recordings. Act-Int: optogenetic response with blue light, neurotransmitter response with redlight, no response with red light plus synaptic blockers, Interluminescence response with red light plus synaptic blockers plus luciferin. Persist-Int: optogenetic response with blue light, Interluminescence response with luciferin. **C** Representative APs for Int (orange) and NT (gray) and **D**, their dV/dt plots. The voltage plotted is the difference in the voltage from the resting membrane potential. Catmull-Rom spline interpolation was used for smoothing. **E** Ladder plots of parameters that were significantly different between all Int APs (Act-Int and Persist-Int) and all NT APs (N = 40 each; Half width (ms): p = 0.0001; After hyperpolarization peak (mV): p < 0.0001). **F** Representative APs elicited through Act-Int and Persist-Int (orange) in comparison to NT-induced (gray) in postsynaptic neurons expressing different excitatory opsins. **G** For recordings of ChR2(C128S)- and CheRiff-expressing neurons, spike frequencies of initial 500 ms after Interluminescent stimulation for Act-Int and Persist-Int are plotted. Spike frequencies for same opsin across Act-Int and Persist-Int are not significantly different (N = 5 per group; ANOVA: ChR2(C128S) Act-Int vs Persist-Int p = 0.7980; CheRiff Act-Int vs Persist-Int p = 0.4799). Spike frequencies between different opsins within Act-Int or Persist-Int are not significantly different (N = 5 recordings per group; ANOVA; Act-Int ChR2(C128S) vs CheRiff p = 0.9710; Persist-Int ChR2(C128S) vs CheRiff p = 0.2492). **H** For recordings of ChR2(C128S)-expressing postsynaptic neurons, change in baseline membrane voltage (ΔmV) for Act-Int (POMC sbGLuc) and Persist-Int (GeNL_SS-ICAM) are not significantly different (N = 13 Act-Int, N = 5 Persist-Int; two-way Welch’s t-test: p = 0.9470).

Cultures were first exposed to blue light to activate opsins directly in striatal postsynaptic neurons (**Fig. 3B**). For Act-Int: red light activated presynaptic ChrimsonR^27^ (postsynaptic ChR2(C128S) is not activated by red light) and neurotransmitter (NT)-mediated postsynaptic responses were recorded (**Fig. 3B** upper panel). NT-mediated postsynaptic activity in response to red light was confirmed by loss of postsynaptic activity in the presence of synaptic blockers and the restoration of postsynaptic responses in response to red light in the presence of luciferin and synaptic blockers isolated Act-Int signaling (**Fig. 3B** upper panel). For Persist-Int: blue light was used to activate postsynaptic opsins and the presence of postsynaptic activity in response to luciferin (without synaptic blockers) confirmed Persist-Int signaling (**Fig. 3B** lower panel).

We compared features of postsynaptic APs mediated by Act-Int and Persist-Int (Int-APs) to control postsynaptic APs generated by neurotransmitters in the same recordings (NT-APs) (**Fig. 3C-E**). Each recording had similar optogenetic responses to blue light and the membrane potentials of postsynaptic neurons were not significantly different across recordings (**Supplementary Fig. S1A, B**; N = 4-13 recordings each; Kruskal-Wallis overall p = 0.0113 (blue light), overall p = 0.4832 (membrane potential); see Table S2 for complete p-values).

We compared postsynaptic responses in cultures mediated by Act-Int and Persist-Int and across neurons expressing ChR2(C128S), CheRiff^23^, ChRmine^24^, and VChR1^28^ (for representative examples of individual APs see **Fig. 3F**). We did not find statistically significant differences in the properties of postsynaptic APs mediated by Act-Int and Persist-Int (see for example **Supplementary Fig. S2A, B**; N = 7-8 recordings each; Kruskal-Wallis; all comparisons p > 0.9999 (half-width), all comparisons p > 0.9999 (Afterhyperpolarization peak)). This includes the rate of rise at the foot of the AP prior to spike threshold consistent with similar postsynaptic processes (**Fig. 3C, D**). We therefore combined these data for comparison to neurotransmitter-mediated postsynaptic APs.

We compared various features of the AP including half-width (at 50% of AP peak), the size of the afterhyperpolarization (AHP), and dV/dt plots. Some parameters were similar between Int-APs and NT-APs including AP latency, threshold, and peak amplitude, but NT-APs had shorter half widths and larger afterhyperpolarizations compared to Int-APs (**Fig. 3E**; N = 43 recordings for all; two tailed Wilcoxon Signed Rank (both); half-width NT vs Int p = 0.0003; AHP NT vs Int p < 0.0001).

While there was considerable variation in spike rates among recordings (**Fig. 3G**), there was no evidence for obvious differences in the frequency of APs during the first 500 ms following Int stimulation with Act-Int versus Persist-Int using two different opsins (ChR2(C128S) and CheRiff).

To further compare Act-Int and Persist-Int we quantified the change in base membrane potential induced by each method by measuring the difference between resting membrane potential (i.e., membrane potential before Int onset) and shifted baseline membrane potential at the saturation level during Int^29,30^ (ΔmV; **Supplementary Fig. S3**). We recorded the change in baseline (in mV) after Int of ChR2(C128S) expressing postsynaptic neurons paired with either POMC sbGLuc (Act-Int) or GeNL_SS-ICAM-Nrxn3b (Persist-Int). No significant difference in postsynaptic activation was observed between the two Int paradigms (**Fig. 3H**; N = 13 recordings (Act-Int) and 5 recordings (Persist-Int); unpaired two-way Welch’s t-test; p = 0.9470).

### Persist-Int synaptic transmission

To assess the robustness of light-mediated synaptic signaling we recorded postsynaptic responses in opsin expressing neurons using different Persist-Int constructs and experimental configurations. We varied the luciferase, the sequences separating luciferase and membrane anchor, and the opsin (**Fig. 4**).

**Fig. 4.**
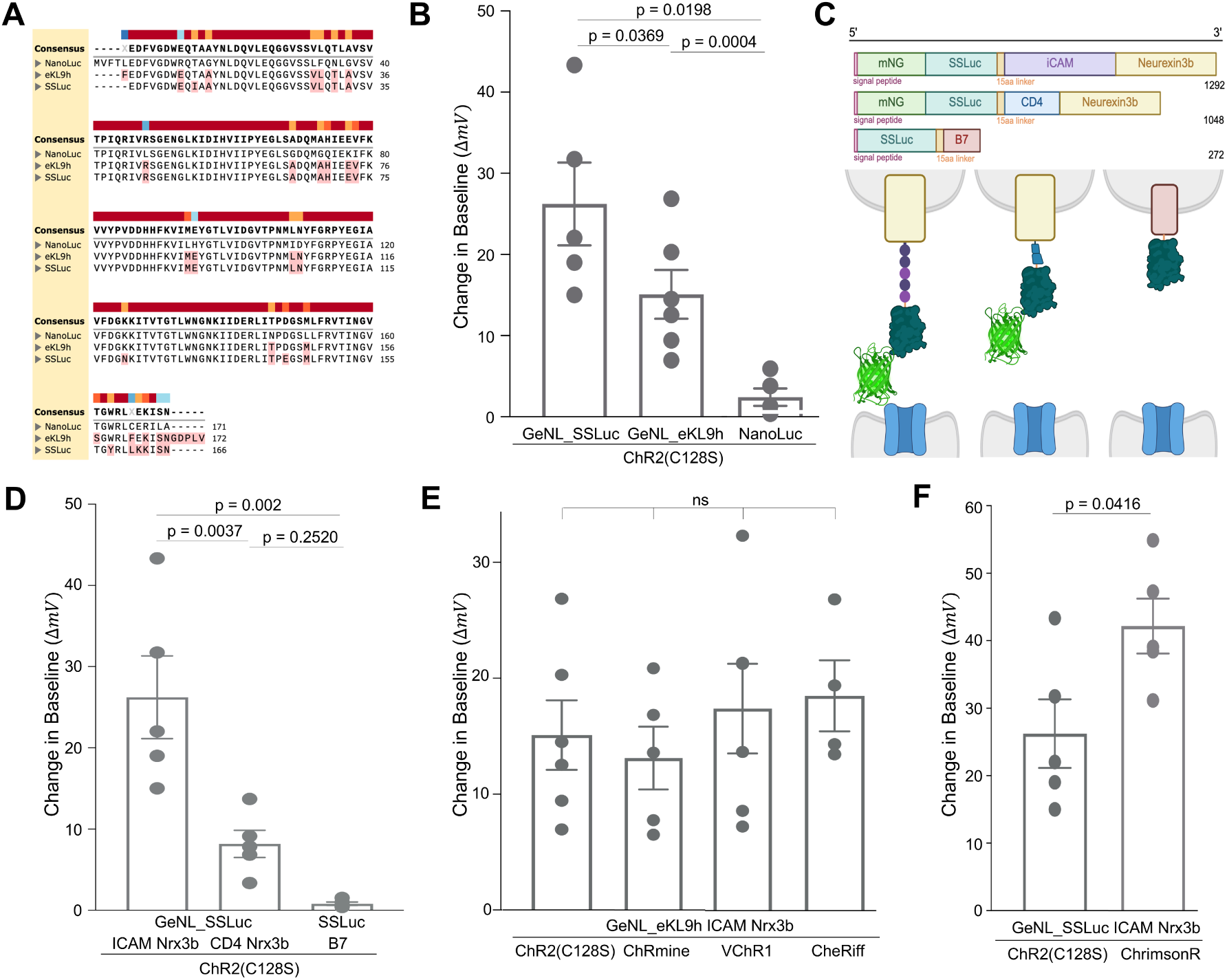
Persist-Int synaptic transmission. **A** Comparison of sequences of NanoLuc, eKL9h, and SSLuc. Mutations were introduced into NanoLuc to yield brighter variants eKL9h and SSLuc. Luciferase brightness is as follows: NanoLuc < eKL9h < SSLuc. **B** Change in baseline membrane voltage in postsynaptic ChR2(C128S)-expressing neurons was compared for different presynaptic Persist-Int constructs with the format of Luciferase-ICAM-Nrx3b. Increasing luciferase brightness was positively correlated to an increase in the change in membrane voltage (N = 5 for GeNL_SS and NanoLuc, N = 6 for GeNL_eKL9h; ANOVA; SSLuc vs eKL9h p = 0.0369; eKL9h vs NanoLuc p = 0.0004; SSLuc vs NanoLuc p = 0.0198). **C** Different sequences were used to tether SSLuc to the extracellular surface of the presynaptic membrane and facing into the synaptic cleft. Sequences differed in length and composition (top) and potentially reached different distances across the synaptic cleft (bottom schematic). **D** The change in baseline membrane voltage in postsynaptic neurons expressing ChR2(C128S) induced by luciferin-mediated light emission from Persist-Int presynaptic constructs. The change in the postsynaptic response in GeNL-SSLuc-ICAM-Nrx3b expressing co-cultures was significantly higher compared to GeNL-SSLuc-CD4-Nrx3b and SSLuc-B7 expressing co-cultures, while postsynaptic responses in GeNL_SS-CD4 and SSLuc-B7 co-cultures were not significantly different (N = 5 recordings each; ANOVA; ICAM vs CD4 p = 0.0037; ICAM vs B7 p = 0.002; CD4 vs B7 p = 0.2520). **E** Change in baseline membrane voltage in postsynaptic ChR2(C128S)-, ChRmine-, VChR1-, and CheRiff-expressing neurons was compared for presynaptic construct GeNL_eKL9h-ICAM-Nrx3b. There was no significant difference between any opsin type upon bioluminescent stimulation (N = 6 recordings (ChR2(C128S)), 5 recordings (ChRmine), 6 recordings (VChR1), 4 recordings (CheRiff); ANOVA; ChR2(C128S) vs ChRmine p = 0.9721; ChR2(C128S) vs VChR1 p = 0.9496; ChR2(C128S) vs CheRiff p = 0.8945; ChRmine vs VChR1 p = 0.7834; VChR1 vs CheRiff p = 0.9958). **F** Change in baseline membrane voltage in postsynaptic ChR2(C128S)- and ChrimsonR-expressing neurons was compared for presynaptic construct GeNL_SS-ICAM-Nrx3b. More sensitive opsin ChrimsonR was found to induce significantly higher increases in membrane voltage (N = 5 recordings each; two-way Welch’s t-test; p = 0.0416).

We compared postsynaptic responses in co-cultures expressing luciferases of different brightness in Persist-Int constructs using the ICAM-Nrxn3b backbone. NanoLuc-ICAM-Nrxn3b, GeNL_eKL9h-ICAM-Nrxn3b, and GeNL_SS-ICAM-Nrxn3b contain luciferases of increasing brightness (NanoLuc < eKL9h < SSLuc). SSLuc and eKL9h are brighter variants of NanoLuc^15^ (**Fig. 4A**). The largest ChR2(C128S) postsynaptic responses induced by light were recorded from GeNL_SS-ICAM-Nrxn3b expressing co-cultures and luciferase brightness was positively correlated with the size of membrane depolarization induced by bioluminescent stimulation (**Fig. 4B**; N = 5 recordings (GeNL_SS and NanoLuc), N = 6 recordings (GeNL_eKL9h); ANOVA; SSLuc vs eKL9h p = 0.0369; eKL9h vs NanoLuc p = 0.0004; SSLuc vs NanoLuc p = 0.0198). This dependence on donor brightness is consistent with a model in which local excitation rate, rather than bulk photon propagation, limits postsynaptic activation.

We also varied the type and length of sequences between luciferase and the membrane targeting protein. The constructs used were GeNL_SS-ICAM-Nrxn3b, GeNL_SS-CD4-Nrxn3b, and SSLuc-B7 and overall construct lengths ranged from 1292 to 272 amino acids (**Fig. 4C**). Whole cell postsynaptic responses in ChR2(C128S) expressing cells were largest for the longest ICAM sequence. Sequence length was positively correlated with the size of the membrane depolarization induced by bioluminescent light stimulation (**Fig. 4D**; N = 5 recordings each; ANOVA; SS_iCAM vs SS_CD4 p = 0.0037; SS_iCAM vs SS_B7 p = 0.0002; SS_CD4 vs SS_B7 p = 0.2520). This effect is consistent with a distance-dependent mechanism in which extending the donor into the synaptic cleft increases the probability of productive donor–opsin interactions.

Finally, we compared light-induced responses using different postsynaptic opsins (ChR2(C128S), ChRmine, VChR1, and CheRiff) paired with GeNL_eKL9h-ICAM-Nrxn3b expressing presynaptic cells. Responses were similar for these constructs (**Fig. 4E**; N = 6 recordings (ChR2(C128S)), 5 recordings (ChRmine), 6 recordings (VChR1), 4 recordings (CheRiff); ANOVA; ChR2(C128S) vs ChRmine p = 0.9721; ChR2(C128S) vs VChR1 p = 0.9496; ChR2(C128S) vs CheRiff p = 0.8945; ChRmine vs VChR1 p = 0.7834; ChRmine vs CheRiff p = 0.7136, VChR1 vs CheRiff p = 0.9958). But, when paired with the bright light emitter GeNL_SS-ICAM-Nrxn3b (**Fig. 4F**) ChrimsonR supported the largest Persist-Int responses when compared to ChR2(C128S) (N = 5 recordings each; two-way Unpaired Welch’s t-test; p = 0.0416).

### Act-Int synaptic transmission

For Act-Int, we similarly compared the efficiency of different construct designs, presynaptic light emitters, and postsynaptic light sensors. In Act-Int, the luciferase is targeted to vesicles via the first 26 amino acids of the human pro-opiomelanocortin prehormone (hPOMC1-26) and released into the synaptic cleft after vesicle fusion (**Fig. 5Ai**). We generated constructs that expose the luciferase to the synaptic cleft but do not release it (**Fig. 5Aii, iii**). We attached the luciferase to the luminal end of a truncated version of synaptobrevin/VAMP2 (TMD only^31^) (**Fig. 5Aii**), and to the luminal C-terminus of the vesicular GABA transporter VGAT^32^ (**Fig. 5Aiii**). These attachments expose the luciferase to the cleft with activity driven vesicle fusion to the presynaptic membrane, but without releasing it. Utilizing neuropeptide tags such as hPOMC1-26 will release luciferases mainly via large dense core vesicles (LDCVs)^16,33^, while vesicular amino acid transporters such as VGAT preferentially target small clear vesicles (SCVs)^34^, and synaptobrevin is present in both vesicle types^35,36^. We found no consistent difference in the efficacy of presynaptic POMC sbGLuc, Synb-sbGLuc, and VGAT-sbGLuc in Act-Int experiments to couple presynaptic depolarization to a postsynaptic opsin-mediated response (**Fig. 5B**; N = 5 recordings (Synb), 5 recordings (VGAT), 13 recordings (hPOMC); Kruskal-Wallis; Synb vs VGAT p = 0.6762; Synb vs hPOMC p > 0.9999; VGAT vs hPOMC p > 0.9999).

**Fig. 5.**
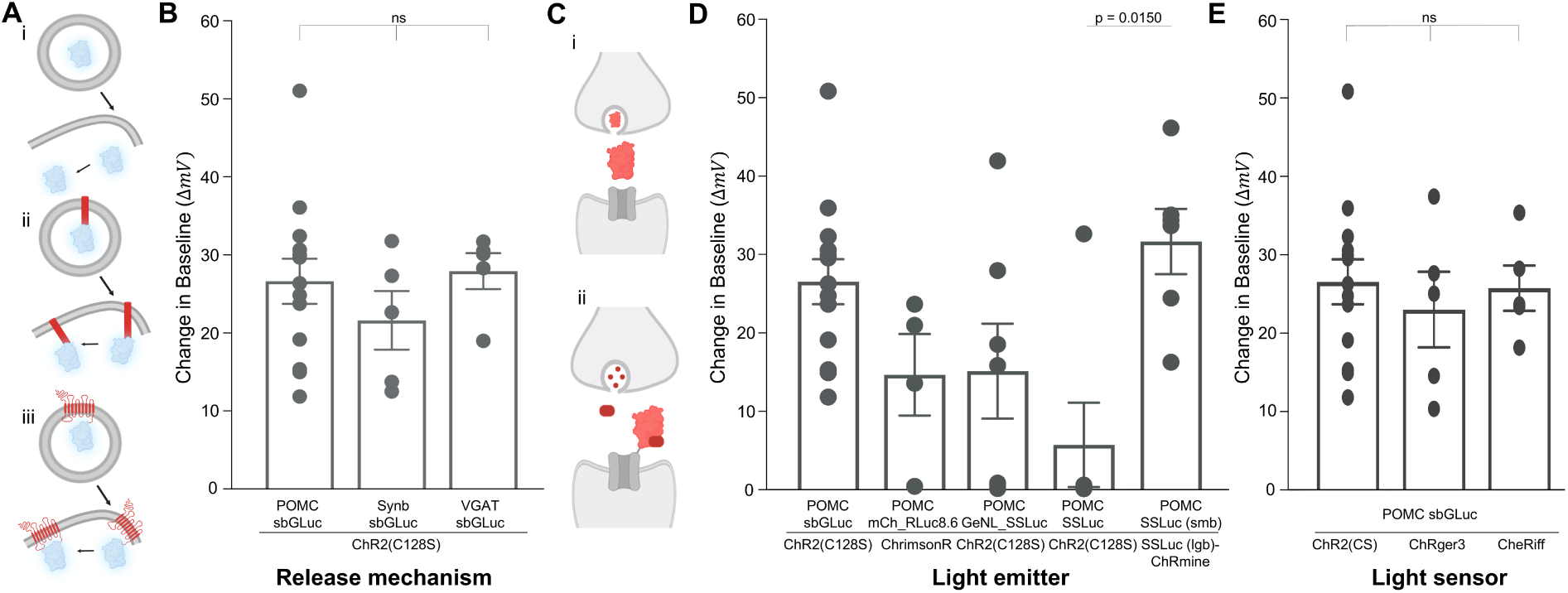
Act-Int synaptic transmission. **A** Different versions of Act-Int were designed using vesicle targeting sequences: (i) the human proopiomelanocortin (hPOMC) signal sequence, (ii) a truncated synaptobrevin2 TMD (Synb), (iii) the vesicular inhibitory amino acid transporter (VGAT). All sequences shuttle the luciferase into synaptic vesicles in presynaptic neurons. Vesicles are released into the synaptic cleft upon presynaptic activation, allowing luciferase to interact with luciferin and produce bioluminescence to activate postsynaptic opsin-expressing neurons. **B** Change in baseline membrane voltage in postsynaptic ChR2(C128S)-expressing neurons was measured for hPOMC sbGLuc, Synb-sbGLuc, and VGAT-sbGLuc Act-Int constructs. All targeting sequences resulted in large changes in membrane voltage upon luciferin application, without significant differences between groups (N = 5 recordings (Synb), 5 recordings (VGAT), 13 recordings (hPOMC); Kruskal-Wallis; Synb vs VGAT p = 0.6762; Synb vs hPOMC p > 0.9999; VGAT vs hPOMC p > 0.9999). **C** Act-Int with intact luciferase (i) versus split luciferase (ii). **D** Change in baseline membrane voltage in postsynaptic opsin-expressing neurons was measured for hPOMC Luc-Act-Int constructs using the intact luciferases sbGLuc, mCherry 22.0_RLuc8.6, GeNL_SS, and SSLuc and the split luciferase SSLuc_smb_. The only significant difference in levels of change in membrane voltage was found for SSLuc that showed decreased responses compared to SSLuc_smb_. All other comparisons were non-significant (N = 13 recordings (sbGLuc), 5 recordings (RLuc), 7 recordings (GeNL_SS), and 6 recordings (SSLuc); Kruskal-Wallis; sbGLuc vs RLuc p = 0.8383; sbGLuc vs GeNL_SS p > 0.9999; sbGLuc vs SSLuc p = 0.0545; RLuc vs GeNL_SS p > 0.9999; RLuc vs SSLuc p > 0.9999; RLuc vs SSLuc_smb_ p = 0.2361; GeNL_SS vs SSLuc p > 0.9999; GeNL_SS vs SSLuc_smb_ p = 0.3826; SSLuc vs SSLuc_smb_ p = 0.0150). **E** Change in baseline membrane voltage in postsynaptic neurons expressing various channelrhodopsins (ChR2(C128S), ChRger3, CheRiff) was assessed for presynaptic Act-Int construct hPOMC sbGLuc. There was no significantly different change in membrane voltage between postsynaptic constructs (N = 13 recordings (ChR2(C128S)), 5 recordings (ChRger3), 4 recordings (CheRiff); ANOVA; ChR2(C128S) vs ChRger3 p = 0.7752; ChR2(C128S) vs CheRiff p = 0.9873; ChRger3 vs CheRiff p = 0.8982).

Like Persist-Int, we also compared the efficiency of different luciferases in Act-Int to couple to postsynaptic activity. Using the hPOMC vesicle targeting sequence, we compared presynaptically released intact luciferases (**Fig. 5Ci**) with split luciferases where a “small bit” (smb) is released from presynaptic vesicles, and the “large bit” (lgb) is tethered to the postsynaptic opsin (**Fig. 5Cii**). We generated Act-Int constructs POMC mCherry 22.0_RLuc8.6^14^, POMC GeNL_SS, POMC SSLuc, and POMC SSLuc_smb_ (amino acids 159-169 of SSLuc) along with the classical POMC sbGLuc and paired them with postsynaptic neurons expressing ChR2(C128S) (for sbGLuc, GeNL_SS, and SSLuc), ChrimsonR (for mCherry 22.0_RLuc8.6), or a lgb (amino acids 1-158 of SSLuc)-ChRmine fusion (for SSLuc_smb_) (**Fig. 5D**). Light emitters in presynaptic neurons were co-expressed with excitatory opsins ChrimsonR (sbGLuc, GeNL_SS, SSLuc and SSLuc_smb_) or ChR2(C128S) (mCherry 22.0_RLuc8.6) to induce presynaptic activation with red or blue light, respectively.

Postsynaptic responses evoked by Act-Int bioluminescence were recorded in 13/13 recordings for sbGLuc, 3/5 for mCherry 22.0_RLuc8.6, 4/7 for GeNL_SS, 1/6 for SSLuc, and 6/6 for SSLuc_smb_. The size of the depolarization induced by these constructs were not consistently different between the presynaptic luciferases sbGLuc, mCherry 22.0_RLuc8.6, GeNL_SS, and SSLuc_smb_, but SSLuc alone (lacking the fluorescent protein mNeonGreen) was less effective compared to SSLuc_smb_ (N = 13 recordings (sbGLuc), 5 recordings (RLuc), 7 recordings (GeNL_SS), and 6 recordings (SSLuc); Kruskal-Wallis; sbGLuc vs RLuc p = 0.8383; sbGLuc vs GeNL_SS p > 0.9999; sbGLuc vs SSLuc p = 0.0545; RLuc vs GeNL_SS p > 0.9999; RLuc vs SSLuc p > 0.9999; RLuc vs SSLuc_smb_ p = 0.2361; GeNL_SS vs SSLuc p > 0.9999; GeNL_SS vs SSLuc_smb_ p = 0.3826; SSLuc vs SSLuc_smb_ p = 0.0150). Luciferin addition without activation of presynaptic neurons did not alter the membrane voltage of the postsynaptic cell (**Supplementary Fig. 4**).

We next compared the efficacy of different postsynaptic channelrhodopsins in co-cultures with POMC sbGLuc expressing presynaptic neurons. Postsynaptic channelrhodopsins ChR2(C128S), ChRger3^22^, or CheRiff, were compared. All postsynaptic light-sensitive proteins induced postsynaptic responses and, within the parameters of our testing, produced comparable responses within the resolution of our measurements (**Fig.5E**; N = 13 recordings (sbGLuc), 5 recordings (ChRger3), 4 recordings (CheRiff); ANOVA; ChR2(C128S) vs ChRger3 p = 0.7752; ChR2(C128S) vs CheRiff p = 0.9873; ChRger3 vs CheRiff p = 0.8982).

### Act-Int efficiency of synaptic transmission

In Act-Int, synaptic transmission depends on activation of the presynaptic neuron. To study the relationship between presynaptic activity and postsynaptic responses, we generated co-cultures of presynaptic neurons expressing the vesicle-targeted luciferase POMC-sbGLuc and the red light sensitive excitatory opsin ChrimsonR, and postsynaptic neurons expressing ChR2(C128S) (**Fig. 6A**). ChR2(C128S) is not activated by red light therefore, red light stimulation only activates presynaptic neurons. Red light of 2 ms to 32 ms in duration triggered postsynaptic responses with increasing numbers of APs as the stimulus duration increased (**Fig. 6B**). The NT response, recorded first, was compared to the Act-Int response within the same neurons in the presence of synaptic blockers. The postsynaptic responses in NT- and Int-mediated synaptic transmission were very similar as the size of the presynaptic stimulation increased (**Fig 6C**; N = 5 recordings (2 ms NT and Int), 3 recordings (4 ms and 16 ms NT and Int), and 4 recordings (32 ms NT and Int); two way Wilcoxon Matched-Pairs Sign Rank (2 ms), two way Paired t-test (4ms, 16ms, 32ms); 2 ms NT vs Int p = 0.7500; 4 ms NT vs Int p = 0.6667, 16 ms NT vs Int p = 0.7745, 32 ms NT vs Int p = 0.9851). The duration of red light activation required to elicit 1, 5, 10, 15, and 20 Int and NT action potentials were basically identical (**Fig. 6D**).

**Fig. 6.**
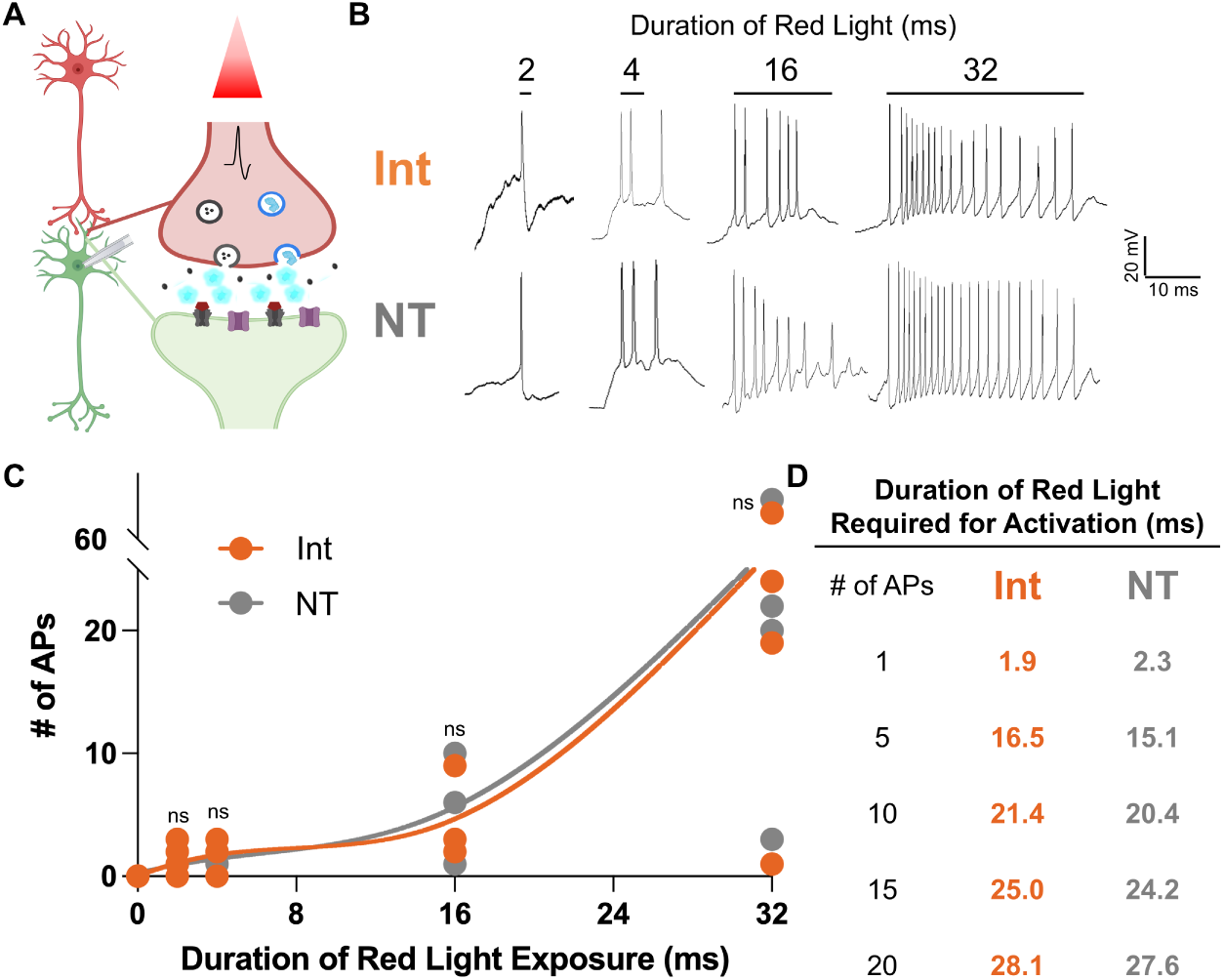
Act-Int efficiency of synaptic transmission. **A** Act-Int schematic: Presynaptic luciferase-expressing neurons were activated by red light, releasing vesicle bound luciferase and neurotransmitter (NT) molecules. To isolate Act-Int effects without NT effects on postsynaptic neurons synaptic blockers (SB) together with the luciferin CTZ was added. **B** Experimental design: Presynaptic neurons were activated by exposure to red light of varying durations (2 – 32 ms) and postsynaptic action potentials (APs) elicited by Act-Int (Int) or by endogenous NT were recorded. **C**. Postsynaptic neurons were recorded after a single exposure to red light of a given length (N = 3 to 5 recordings for each light exposure time, each data point is one neuron, NT and Int data for each respective time point are from same neuron). Restricted cubic smoothing spline with 2000 points was used to interpolate data. No significant differences in the resulting number of action potentials between Int and NT recordings was found at each time point (two way Paired Wilcoxon Matched-Pairs Sign Rank Test: 2ms p = 0.5716; two way Paired t-test; 4 ms p = 0.6667, 16 ms p = 0.7745, 32 ms p = 0.9851). **D** Using interpolation data points of smoothing spline, the durations of red light activation required to elicit 1, 5, 10, 15, and 20 Int and NT action potentials were identified.

### Synapse specificity of Interluminescence transmission

While designed to modify synaptic transmission, we wanted to experimentally test whether Interluminescence effects are restricted to synapses. For Act-Int, it is possible that luciferases released during Act-Int protocols diffuse away from the synapse and influence postsynaptic responses in neighboring, not synaptically connected opsin-expressing neurons. We recorded from ChR2(C128S) opsin-expressing neurons synaptically connected to luciferase-expressing presynaptic neurons (POMC sbGLuc P2A ChrimsonR dTomato), and from nearby neurons that were not synaptically connected (**Fig. 7A**). “Synaptically connected” neurons were identified by red light activation of presynaptic neurons eliciting robust firing of postsynaptic APs and “non-synaptically connected” neurons nearby by the lack of responses. Nearby neurons were in the same 40x field of view (max 0.66 mm apart; **Supplementary Fig. 6**). To confirm that neurons expressed similar levels of opsin, we recorded neuronal responses to blue light (**Fig. 7B**). To establish if two neurons were synaptically connected we exposed the culture to red light (**Fig. 7C**).

**Fig. 7.**
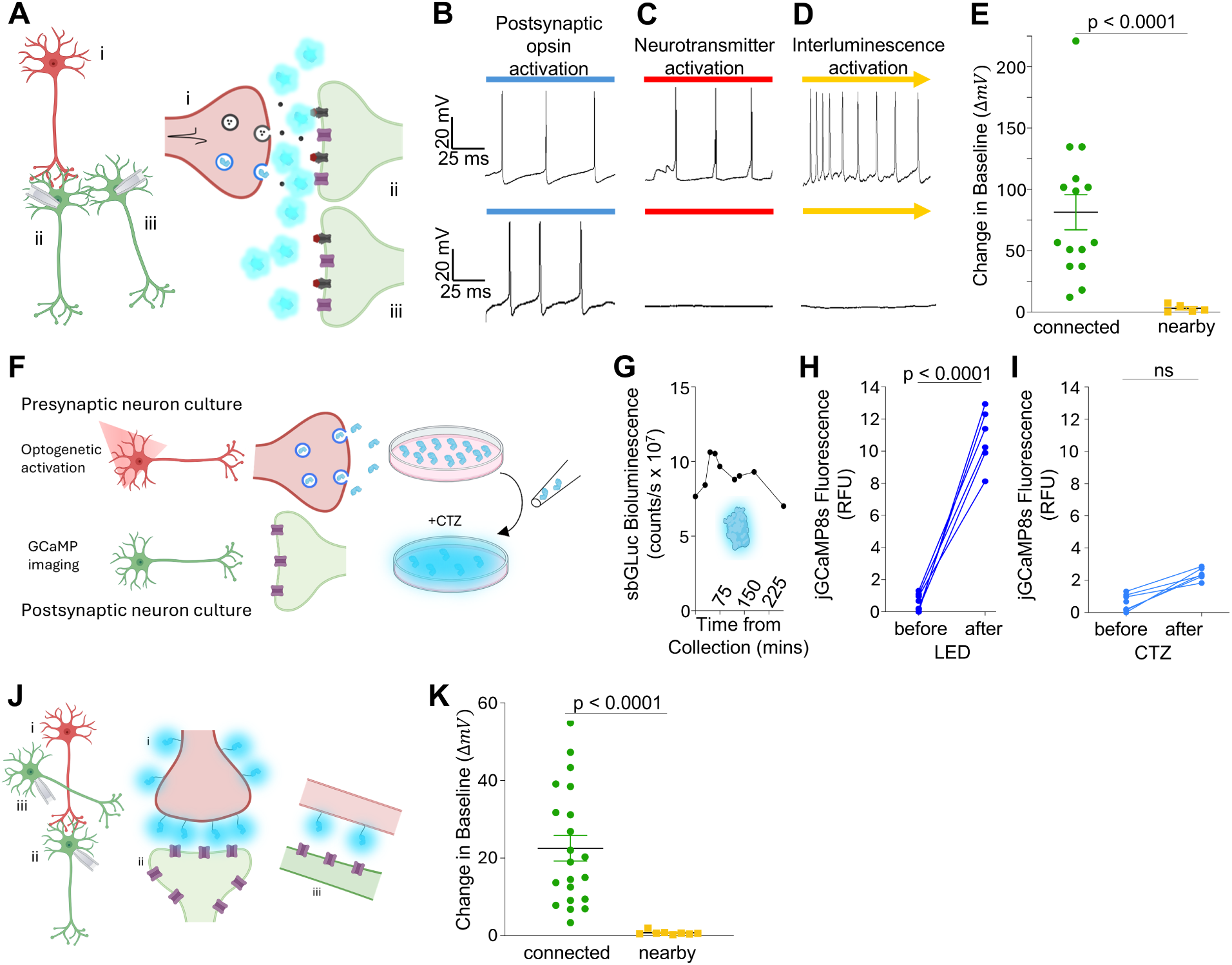
Specificity of Int synaptic transmission. A-I Act-Int specificity of synaptic transmission. **A** Experimental design: Postsynaptic opsin expressing neurons were identified by their green fluorescence and patched. Neurons were either synaptically connected to luciferase and ChrimsonR expressing neurons or not. **B** Recording from opsin expressing neurons under blue light (480 nm) exposure revealed their level of opsin expression. **C** Recording from opsin expressing neurons under red light (580 nm) exposure showed the postsynaptic neuron’s neurotransmitter response to synaptically connected (upper trace) versus non-connected (lower trace) presynaptic neurons. **D** Recording from opsin expressing neurons under red light exposure in the presence of synaptic blockers and CTZ allows isolation of the postsynaptic neuron’s response to bioluminescence-mediated opsin activation. A response is elicited only in the synaptically connected (upper trace) but not in the nearby non-connected (lower trace) neuron. **E** Summary of recordings conducted in the presence of synaptic blockers and exposure to red light (1 s) showing the change in baseline membrane potential when CTZ was added to the recording bath. Changes in membrane potential are only seen in synaptically connected neurons in the presence of CTZ (N = 13 neurons (connected) and 5 neurons (nearby); two-way Unpaired Welch’s t-test p < 0.0001). **F** Schematic of conditioned media transfer experiment. Cultures of neurons expressing presynaptic construct hPOMC sbGLuc P2A ChrimsonR dTom were stimulated with continuous red light and media containing vesicle released sbGLuc was collected and added to separate cultures of neurons expressing postsynaptic construct ChR2(C128S) as well as jGCaMP8s, luciferin was added and change in jGCaMP8s fluorescence as a proxy for neuronal activation was measured. **G** Presynaptic media containing sbGLuc was removed from cultures and bioluminescence was measured at different time points from collection, demonstrating stability of the luciferase. **H** Control cultures of postsynaptic opsin expressing neurons were stimulated with blue LED (470 nm) to test jGCaMP8s fluorescence in response to standard optogenetic activation. There was a significant increase in fluorescence compared to baseline (N = 6 wells; Kruskal-Wallis; p < 0.0001). **I** jGCaMP8s fluorescence of wells with postsynaptic opsin expressing neurons given presynaptic sbGLuc containing media and CTZ was measured and compared to baseline. There was no significant increase in fluorescence (N = 6 wells; Kruskal-Wallis; p = 0.2438). **J-K Persist-Int specificity of synaptic transmission. J** Design of neuron experiments: Postsynaptic opsin expressing neurons were identified by their green fluorescence and patched. Neurons were either synaptically connected to luciferase expressing neurons or were unconnected nearby neurons in the same field of view. **K** Summary of recordings showing the change in baseline membrane potential when luciferin (hCTZ) was added to the recording bath. Recordings from 5 combinations of Persist-Int (presynaptic) and opsin (postsynaptic) are shown: GeNL_eKL9h-ICAM-Nrxn3b x ChR2(C128S), GeNL_SS-ICAM-Nrxn3b x ChR2(C128S) and ChrimsonR, GeNL_SS-CD4-Nrxn3b x ChR2(C128S) and ChRmine. Changes in membrane potential are only seen in synaptically connected neurons (N = 20 neurons (connected) and 8 neurons (nearby); two-way Unpaired Welch’s t-test p < 0.0001).

Red light activates the opsin in presynaptic neurons, resulting in the release of neurotransmitters and luciferase, but it does not activate postsynaptic opsins directly. As expected, only the synaptically connected postsynaptic neurons responded with AP firing. To separate Interluminescence from the endogenous neurotransmitter action we repeated red light stimulation in the presence of synaptic blockers and CTZ (**Fig. 7D**). Bioluminescence activation of postsynaptic opsins occurred only in neurons synaptically connected to the luciferase-expressing presynaptic neuron, whereas a nearby neuron not synaptically connected did not respond. **Figure 7E** summarizes the changes in membrane potential seen across recordings. Throughout all recordings, only neurons that had a response to red light induced presynaptic neuron firing also showed the Act-Int response (N = 13 recordings (connected) and 5 recordings (nearby); two-way Unpaired Welch’s t-test p < 0.0001), strongly suggesting that productive activation occurs predominantly at synaptic contacts rather than through spillover to nearby cells.

In a related but different experiment, we established separate cultures of presynaptic and postsynaptic neurons; presynaptic neurons expressed POMC sbGLuc P2A ChrimsonR dTomato and postsynaptic neurons expressed ChR2(C128S) and the fluorescent Ca^2+^ indicator jGCaMP8s^37^. Optogenetic stimulation of the presynaptic culture with red light for 5 sec released luciferase into the medium. Luciferase-containing medium was then transferred to postsynaptic cultures. After a one-hour rest period, CTZ was added to the postsynaptic cultures and neuronal activation was assessed by measuring change in Ca^2+^ dependent fluorescence over baseline (**Fig. 7F**). To ensure that the luciferases released into the medium were functional, we showed bioluminescence emission of the transfer solution at times after collection (**Fig. 7G**). The luciferase in the transfer solution was functional for several hours after collection. To control for opsin function and GCaMP expression, we exposed postsynaptic neurons to blue light for 2 mins and observed increased activity. By contrast, postsynaptic cultures exposed to bioluminescence in the medium did not exhibit increased neuronal activity (**Fig. 7H, I**; N = 6 wells each; Kruskal-Wallis; LED before vs LED after p < 0.0001; CTZ before vs CTZ after p = 0.2438). These data further support the model in which synaptic transmission through Act-Int is a local signaling event that does not spill over to affect nearby neurons.

For Persist-Int, the luciferases are tethered to the membrane and thus do not leave the synaptic cleft. However, despite the Nrxn3b sequence directing the luciferase to the presynaptic membrane, it is likely that at least to some extent luciferases will be distributed across the surface of the presynaptic neuron, opening the possibility of Interluminescence activation of nearby, not synaptically connected opsin-expressing neurons (**Supplementary Fig. 6**). To experimentally address this question, we conducted recordings from synaptically connected and nearby neurons analogous to those described above for Act-Int (**Fig. 7J**). In mixed cultures of cortical Persist-Int expressing neurons and striatal opsin expressing neurons Interluminescence effects were seen only in striatal neurons synaptically connected to cortical neurons, but not in nearby neurons not synaptically connected (**Fig. 7K**; N = 20 recordings (connected) and 8 recordings (nearby); p < 0.0001), again strongly suggesting that opsin activation by luciferases extended from presynaptic neurons occurs predominantly at synaptic contacts.

## Discussion

Interluminescence can achieve signal transmission at synapses independent of the action of a neurotransmitter. This type of ‘optical synapse’ can be activity-dependent (‘Act-Int’) where genetically encoded luciferase is targeted to synaptic vesicles and released along with neurotransmitter in response to presynaptic activation. Depending on the nature of the genetically encoded postsynaptic opsin, Act-Int can be excitatory or inhibitory. ‘Persist-Int’ similarly depends on expressing genetically encoded luciferase in presynaptic neurons and opsin in postsynaptic cells, but in this case synaptic signaling is activity independent, as luciferase is tethered to the extracellular surface of the presynaptic membrane and light emission depends only on the presence of luciferin.

Here we describe features of Act-Int and Persist-Int and show creation of novel synaptic signaling pathways in cultured neurons. Central to predicting how optical synapses can be used to modulate existing synaptic connections and constitute synaptic signaling *de novo* is showing how Act-Int and Persist-Int are similar and/or different from neurotransmitter-mediated transmission. To that end, we evaluated the effectiveness of Act-Int and Persist-Int and the properties of postsynaptic responses relative to neurotransmitter-induced synaptic transmission in the same cells.

Aside from longer duration of Int-mediated APs we did not observe any obvious differences when comparing NT-transmission and Int-transmission. Further, we did not find evidence for differences between Act-Int and Persist-Int regarding AP features, spike frequencies and change in base membrane potential. However, further in-depth analysis of larger numbers of recordings might reveal subtle differences.

The key molecules that mediate Int can be modified depending on the desired level of control and outcome. For both Act-Int and Persist-Int, a variety of luciferases and opsins can be employed, following the general rule that, not surprisingly, the brightest light emitter, GeNL_SS, paired with the most sensitive opsin tested, ChrimsonR, generated the largest postsynaptic response. Aside from these shared attributes, each Int method has specific parameters amenable to tuning.

For Persist-Int, the luciferase is tethered to the presynaptic membrane and thus distance to the postsynaptic opsin is dependent on the length of sequence between the presynaptic membrane anchor and the luciferase. We showed that altering the length of this sequence affected the efficacy of Persist-Int. For the GeNL-eKL9h-ICAM-Nrxn3b construct, the length of sequence extending into the synaptic cleft after the neurexin3b transmembrane anchor is the 15 amino acid linker (5.25.nm) (3.5 angstrom per AA^38^) + ICAM (18.7 nm^39^)+ GeNL-eKL9h spheroid diameter^40^ (4.7 nm), totaling 28.7 nm. Given an average synapse length of 20 – 30 nm^41^ the luciferase could extend across much of the synaptic cleft, bringing it into a range favorable for efficient FRET to the opsin. However, these are theoretical calculations, and the actual geometry will be influenced by linker flexibility, molecular orientation, and folding properties of the fusion protein. In addition, while Nrxn3b is expected to enrich the construct at presynaptic specializations, we do not assume that expression is confined exclusively to boutons. Rather, our results suggest that functional specificity arises from the strong distance dependence of luciferase-opsin coupling.

For Act-Int, the luciferase is released and can be assumed to cross the synaptic cleft and activate opsins through direct energy transfer mechanisms. Efficacy of the postsynaptic response might be influenced by the number of luciferases released. Considering size/volume of luciferase^40,42^, loading capacity of presynaptic vesicles^43,44^, and number of vesicles present in a mammalian synapse^44,45^, a maximum 5 x 10^5^-16 x 10^5^ molecules of sbGLuc can be released from a single synapse in a one-second red light stimulation of the presynaptic neuron (see **Supplementary Text** for step-by-step calculations). This compares to 16×10^5^-42×10^5^ molecules of glutamate being released in a single one-second red light stimulation of the presynaptic neuron^46^. Again, these are theoretical calculations; for example, these values assume that a one second stimulation induces the release of all available vesicles from the presynaptic neuron and that each vesicle is filled completely with luciferase molecules.

Interestingly, we found similar efficacies in eliciting postsynaptic responses with Act-Int strategies that employ luciferases exposed upon vesicle fusion but stay tethered to the synaptic vesicle rather than being released (vesicle targeting sequences from synaptobrevin and vesicular glutamate transporter versus pro-opiomelanocortin). This opens the possibility of targeting different secretory organelles, large dense core vesicles (POMC), small clear vesicles (VGAT), or both (Synb), taking advantage of different kinetic properties of activity-dependent presynaptic release of luciferase.

We found similar performance in eliciting postsynaptic responses with luciferases from *Gaussia* (sbGLuc), *Oplophorus* (SSLuc), and *Renilla* (RLuc8.6). With red-shifted emission wavelength and luciferin specificity distinct from *Gaussia* and *Oplophorus* based luciferases, the mCherry 22.0_RLuc8.6 construct offers the prospect of multiplexing pre-and postsynaptic luciferase-opsin pairs for differential synaptic manipulation in the same experimental system.

In addition to brightness of the light emitter, a factor to be considered for Act-Int is the stability of the luciferase in the acid environment of the vesicle. Evidence for this is the correlation of pH sensitivity with performance, with the highly pH sensitive SSLuc performing poorly compared to the more pH resistant sbGLuc and RLuc8.6^47,48^ (**Supplementary Fig. 5**). The pH tolerance of SSLuc seems vastly improved when only a small C-terminal peptide of the luciferase, SSLuc_smb_, is packaged in vesicles and released to reconstitute the larger N-terminal part of SSLuc, SSLuc_lgb_, tethered to the postsynaptic opsin. Additional advantages of this split luciferase design are the increased number of small bits that fit into a vesicle compared to intact luciferases, and the increased kinetics of on/offset of bioluminescence due to the rapid removal of the small bit from the large bit.

Both Act-Int optical and NT-mediated chemical synaptic transmission require activity of the presynaptic neuron. We found a perfect overlap between the input-output curves of presynaptic activation and postsynaptic action potential firing generated either through neurotransmitters or through bioluminescence. Thus, Act-Int can be integrated into the activity of endogenous circuits.

The effect of activation of presynaptic neurons on postsynaptic responses can occur through neurotransmitters and through neuropeptides, both released upon presynaptic neuron firing. While neurotransmitter effects are confined to synaptically connected neurons, neuropeptides can have volumetric effects that extend beyond the synaptic cleft to nearby, non-synaptically-connected neurons that express the appropriate receptors. In our experiments, Act-Int behaved more like neurotransmitter-mediated signaling, with bioluminescence driving activity only in opsin-expressing neurons that were synaptically connected to luciferase-expressing neurons. Similarly, Persist-Int exerted its effects on synaptically connected neurons rather than on nearby neurons expressing opsins. For Persist-Int, this functional restriction is important because membrane-tethered luciferase may extend beyond presynaptic boutons, but our same-field paired recordings showed responses in connected neurons and not in nearby non-connected neurons. Thus, although we cannot exclude effects at closely apposed specialized contacts such as en-passant synapses, our results indicate that for both forms of Interluminescence, the dominant effect on postsynaptic neurons arises from synaptically connected presynaptic partners. This makes Interluminescence a highly specific tool for experimenter-controlled synaptic transmission.

The underlying mechanisms of either version of optical synapses can be assumed to involve photon transmission, energy transfer between the chromophores of light emitter and light sensor, or a combination of both. Activation of opsins through Förster resonance energy transfer (FRET), the most efficient mechanism, is inversely proportional to the sixth power of the distance between the luciferase donor and opsin acceptor, requiring a practical distance of 1 – 10 nm^49,50^. In addition to distance, efficiency of FRET is also dependent on the overlap in donor emission and acceptor absorption spectrum, their relative dipole orientation, and the quantum yield of the donor. Thus, effective postsynaptic activation is expected to depend strongly on close apposition of luciferase and opsin, whereas bioluminescence emitted from other parts of the presynaptic neuron is unlikely to contribute substantially to productive signaling at the synapse. For this reason, direct imaging of the overall spatial pattern of bioluminescence in neurons, while informative in principle, would not by itself resolve the functionally relevant component of Interluminescence at the level of individual synaptic contacts. We therefore focused on functional tests of synapse specificity and on experimentally exploring the physical parameters governing luciferase – opsin interaction for Interluminescence.

The order-of-magnitude analysis in **Supplementary Tables 1 and 2** indicates that isotropic radiative transfer from individual luciferase molecules is insufficient to explain opsin activation under standard irradiance benchmarks. This apparent discrepancy is resolved by the observation that Persist-Int signaling is strongly restricted to synaptic contacts and is sensitive to both donor brightness and cleft-spanning architecture. Together, these findings argue that effective transmission depends on nanoscale proximity and molecular organization that increase the local excitation rate experienced by opsins, rather than on bulk photon flux. In this framework, the relevant quantity is the local excitation probability at apposed donor–opsin pairs, which can be substantially higher than predicted by a free-space emission model.

Three other engineered synthetic neurotransmission approaches have previously been introduced. Similar to Act-Int, PhAST (Photon Assisted Synaptic Transmission) utilizes bioluminescence emitted from activated presynaptic neurons to drive optogenetic channels in postsynaptic neurons^51^. Increasing Ca^2+^ concentrations correlating with presynaptic activity induce photon emission from Ca^2+^-dependent luciferases expressed in presynaptic neurons. PhAST has been employed in the model organism *C. elegans* to overcome the behavioral defect from disconnected functional neurotransmission between a sensory neuron and its cognate postsynaptic interneurons, to suppress the endogenous pain response, and to rewire a circuit mediating attractive behavior into an avoidance response^51^. Both types of optical synapses, the Int strategy as well as the PhAST strategy, require the addition of exogenous luciferin to engage the system. The clear advantage of this is experimenter-control over when the engineered synapse is operational. This contrasts with two other systems of synthetic synapses that are based on artificial neuropeptide signaling pathways, where connections cannot be enabled or disabled by the experimenter and persevere throughout the host organism’s entire lifetime. The HySyn (Hydra-derived Synthetic Synapse) system employs heterologous production of a cnidarian neuropeptide through a presynaptic carrier molecule and postsynaptic expression of a Ca^2+^ channel that is also the cognate receptor for the neuropeptide, exploiting the specificity of this neuropeptide–receptor pair to build a synthetic synapse unresponsive to endogenous neuropeptides^52^. The effectiveness of HySyn was confirmed in both *C. elegans* and cultured cells. A similar principle was utilized in rats, where viscerosensory neurons expressing the insect neuropeptide allatostatin were paired with neurons in the brainstem nucleus of the solitary tract expressing allatostatin receptors linked to G protein-coupled inward-rectifier potassium channels^53^. This induced *in vivo* neuropeptide-driven postsynaptic membrane potential inhibition and modulation of blood pressure.

Interluminescence Act-Int and Persist-Int strategies are synapse-specific transmission methods that can be used to achieve transsynaptic communication that is either activity-dependent or activity-independent. The modularity of this method allows expansion into combining the growing arsenal of potent luciferases with the ever-increasing toolset of optogenetic elements, affording experiment-specific tuning of approaches. Each version of Int offers unique features for applications. Act-Int is based on the presynaptic neuron’s activity and thus is embedded in the brain’s endogenous circuit dynamics, synthetically altering the dynamics and flow of internally generated neuronal activity, while Persist-Int can override any status of the presynaptic component, either activity or inactivity. Both modalities allow a new regime of experiments for changing or re-routing information flow. Ongoing effects of enhancing or inhibiting postsynaptic activity can either be augmented or counteracted. These capabilities for changing circuit dynamics can be exploited in basic research for testing specific hypotheses about neural communication, including regarding neural development and plasticity. Int may also provide a unique approach to treating diseases that result from failed signaling in the brain or spinal cord. The increased specificity of targeted pre-to postsynaptic communication will help avoid broad effects on competing target populations that often suffer from canceling each other out.

Our results establish Interluminescence as a versatile and modular platform providing synapse-specific transmission strategies that can be used to achieve transsynaptic communication between partners of choice that is either activity-dependent or activity-independent. The developed toolset provides a new means of interrogating the brain and uniquely contributes solutions not possible with current technologies.

## Materials and Methods

### Plasmids

Coding sequences were cloned into pcDNA3.1 using standard procedures including restriction enzymes and ligation, Gibson cloning, PCR amplification, and gene synthesis. Supplementary Table 3 lists all plasmids used in these studies.

### Luciferin storage and concentrations

Coelenterazine (CTZ) (NanoLight Technology; Pinetop, AZ; NanoLight # 303) and Coelenterazine h (hCTZ) (NanoLight Technology; Pinetop, AZ; NanoLight # 301) were stored in 50 mM stock solutions in NanoFuel solvent (NanoLight # 399). For MEAs recording, hCTZ was applied at 10 μM from a 1 mM working solution of hCTZ in culture media. The NanoFuel vehicle was similarly diluted. For whole cell recordings, hCTZ was applied at 100 μM from a 0.5 mM working solution diluted in extracellular solution.

### Primary Neurons

Male and female embryonic day 18 (E18) rat embryo cortex and striatum samples were obtained from BrainBits, LLC, and processed according to the BrainBits protocol to isolate primary neurons. Briefly, tissue samples were incubated in Hibernate E (without B27 supplement or calcium; HEB, BrainBits) freshly supplemented with 2 mg/ml papain (BrainBits) for 10 min at 30°C. HEB with papain media was replaced by plain HEB medium, then the tissue triturated with a 9” sterile silanized glass Pasteur pipette (BrainBits) for about 1 min until ∼90% tissue dispersal was achieved. Undispersed tissue was allowed to settle to the bottom of the tube (∼1 min) and the solution transferred to a sterile 15 mL tube. The dissociation preparations were centrifuged at 1500 rpm for 10 mins, the supernatant removed, and the pelleted cells resuspended in 200 μL of a pH equilibrated and pre-warmed NbActiv1 medium (BrainBits) and cell density counts obtained.

### Nuclear transfection

E18 primary rat neurons were used for nuclear transfection according to manufacturer’s instructions (Amaxa Rat Neuron Nucleofector Kit # VPG-1003). 1×10^6^ primary neurons were removed from the cell suspension, centrifuged at 1,200 rpm for 7 min at RT °C in a 1.5-mL Eppendorf tube, the cell pellet resuspended in 100 μl Nucleofector Solution, 1 μg plasmid DNA added, and the solution transferred to a cuvette for nuclear transfection (Nucleofector 2b Device (LONZA # AAB-1001); Program G-013). Immediately after nuclear transfection, cells were diluted in NbActiv1 medium, spun to pellet, and resuspended in neuronal plating media (see below).

### Neuronal co-cultures

Co-cultures of cortical neurons transfected with the luciferase construct and cortical or striatal neurons transfected with opsin constructs were mixed in a ratio of 1:1. Cells were grown in Neurobasal Medium (Gibco # 21103-049) supplemented with 5% Fetal Calf Serum (FCS), B-27 supplement (Gibco # 17504-044), and 2 mM Glutamax (Gibco # 35050-061) (NB-FCS). For MEA recordings: 1 × 10^5^ cells per 10 μl NB-FCS per MEA well were plated on the electrode area (60MEA200/30iR-Ti; Multi Channel Systems, Germany), which had been coated with 50 μg/ml laminin (Gibco # 23017-015) and 0.1% polyethyleneimine (Sigma # P3143); for whole cell recordings: 1 × 10^5^ cells per 1 ml NB-FCS were plated on laminin-coated (50 ug/mL) poly-D-lysine coated (15mm, Neuvitro GG-15-PDL) coverslips per well of 12-well tissue culture plates. The following morning, NB-FCS solution was replaced with serum-free medium (NB-Plain medium) in both MEA and 12-well cultures. Thereafter, for MEAs, 50% of the culture medium was replaced with fresh NB-Plain medium every 3–4 days; for whole cell cultures, 300 μL of media was replaced with 500 μL prewarmed and pH equilibrated Neurobasal Plus (NB+) media with B-27 Plus supplement (Gibco # A3582801) at day 3 and then weekly.

### MEA recordings

All MEA recordings were conducted with a MEA2100-Lite-System (Multichannel Systems, Germany) on co-cultures at DIV 14-25. Cultures that did not exhibit spontaneous activity and consistent spiking were not used. All-trans retinal (R2500; Sigma-Aldrich, St. Louis, MO) at a final concentration of 1 μM was added to the culture medium prior to recording. All solutions were pre-warmed to 37°C and the MEA2100 head stage was maintained at 37°C including prior to recording. MEAs were equilibrated for 5–10 minutes after transfer from the CO_2_ incubator. Light stimulation of MEA cultures was achieved with a fluorescent light source of different wavelengths through the objective (Zeiss Observer 1). Pre-warmed NB-Plain media was used to refresh MEAs before returning to the incubator for multiple day recordings. MEA data were acquired using the MC Rack Software, sampled at 10,000 Hz, and analysis performed offline with NeuroExplorer (RRID: SCR_001818) and MC Rack software (Multichannel Systems; RRID: SCR_014955).

### Whole cell recordings

Whole cell current clamp recordings were obtained from primary neurons DIV 14-21 for Persist-Int and DIV 21-28 for Act-Int experiments. Recordings were made from co-culture coverslips attached to the bottom of a recording chamber (RC26-GLP, Warner Instruments) fixed to a microscope stage (BX51WI, Olympus). Cultures were perfused with external buffer (at 1.5 mL/min) containing (in mM): 121 NaCl, 26 NaHCO_3_, 1 NaH_2_PO_4_, 15 D-glucose, and 2.8 KCl, (∼280-295 mOsm/kg) maintained at 37°C ± 1°C and pH buffered using carbogen saturated solution. External buffer was supplemented with 0.2 mM MgCl_2_*6H2O and 0.2 mM CaCl_2_.

Borosilicate glass was used to pull micropipettes (PC-100, Narishige) which had resistances of 2-6 MΩ when filled with intracellular solution (in mM): 8 KCl, 130 K-gluconate, 5 Na_2_-phosphocreatine, 15 HEPES, 0.3 Na_2_-GTP, 4 Na_2_-ATP at pH 7.25, and adjusted to ∼290-300 mOsm/L. The expression of luciferase (presynaptic, red or green) and opsin (postsynaptic, green or red) were identified with epifluorescence. The metal halide light source (130 W, U-HGLGPS, Olympus) had filter cubes for blue (Ex/Em: 480/530 nm, U-MNIBA3, Olympus), green (Ex/Em: 540/600, U-MWIGA3, Olympus) and red (Ex/Em: 580-630/645-695, ET-Alexa633 Filter Set 605/50x, 670/50M w/BX2 cube) excitation. Once a recording was established, blue light was applied through the objective (LUMPLFLN40XW, 433 NA 0.8, Olympus) controlled by an electronic shutter (Lambda SC, Sutter Instruments) to apply specific timed exposures. Maximum light intensity for each color was measured at (in mW/cm2) 16.8 (blue), 36.5 (green), and 39.9 (red) with a light meter (ThorLabs). Solutions were added to the recording chamber via an inlet tube for a final 1:5 dilution.

### jGCaMP8s fluorescence readings and sbGLuc signal experiments

E18 striatal neurons were nucleofected with equal quantities (1 *μ*g each) of pcDNA CAG jGCaMP8s and pcDNA CAG ChR2(C128S)-dTomato and seeded at ∼6×10^4^ cells per well on a poly-d-lysine coated (0.1 mg/mL) black 96-well plate (Greiner Bio-One CAT: 655090) (“postsynaptic” cultures). Concurrently, E18 cortical neurons were nucleofected with 1 *μ*g of pcDNA CAG POMC sbGLuc P2A ChrimsonR dTomato and seeded at ∼2×10^5^ cells per well of a poly-d-lysine coated (0.1 mg/mL) 4-well plate (“presynaptic” cultures). Both cultures were maintained for three weeks with weekly partial media changes (30 μL removed and 50 μL added) before experiments were conducted. At DIV 21, presynaptic cultures were stimulated with 5s of red light (130W, U-HGLGPS, Olympus, Ex/Em: 580-630/645-695 ET-Alexa633 Filter Set 605/50x, 670/50M w/BX2 cube) per well. Immediately following, conditioned media from all wells of presynaptic cultures was collected. One hundred microliters per well of presynaptic media was transferred to a white 96-well plate (Greiner Bio-One CAT: 655098). At various time points post-media collection, CTZ (at final concentration 100 *μ*M) was added to a well and resulting bioluminescent output read on a plate reader (Tecan Spark). In between readings, the conditioned media plate was kept in the incubator until final reading. Remaining conditioned media was added at increasing amounts to postsynaptic cultures; 0% media replacement, 12.5% media replacement, 25% media replacement, 50% media replacement, 75% media replacement, and 100% media replacement, and cultures returned to the incubator to rest for 1 hour. After the rest period, 1 *μ*L of CTZ (final concentration 500 *μ*M) was added to every well except 0% replacement wells and green fluorescence of jGCaMP8s was measured for all wells (including 0% replacement wells) after ∼6 minutes of CTZ exposure on a plate reader with real-time cytometry capabilities (Tecan Cyto100). Upon completion of fluorescence measurements, 0% replacement wells were exposed to 2 minutes of blue LED light (480 nm) to optogenetically stimulate ChR2(C128S) and green fluorescence measurements retaken as a positive control.

### Synaptic blockers

For Act-Int experiments, the following deionized water stock solutions of synaptic blockers (SB) were kept in aliquots at −80°C: D-AP5 (50 μM, Sigma Aldrich # S106), NBQX (10 μM, abcam # ab120046), CGP55845 (100 μM, Sigma Aldrich # SML0594), Gabazine (100 μM, Sigma Aldrich # S106), and Strychnine (1 μM, Sigma Aldrich # S0532). For CTZ and vehicle recordings, CTZ or vehicle stocks were diluted with the SB cocktail to attain a final CTZ concentration of 10 μM or an equivalent volume in case of the vehicle. Final concentration of synaptic blockers when entering recording chamber were 10 *μ*M of D-AP5, 1 *μ*M of NBQX, 20 *μ*M of CGP55845, 20 *μ*M of Gabazine, and 0.2 *μ*M of Strychnine.

### Data analysis and statistics

All analyses were performed with Prism software including running appropriate statistical tests (GraphPad 8.2.1; San Diego, CA). Prism was also used to assess normality of the data to help determine which test to use.

For MEA recordings, data was collected from multiple recordings per plate across several days and across at least 3 separate plates for each type of co-culture (pre-synaptic Persist-Int with postsynaptic excitatory or inhibitory opsin, respectively). Data obtained under the same experimental conditions (blue light, hCTZ, vehicle) were pooled, with N reflecting the number of individual electrodes analyzed across all experiments. The response to direct blue light stimulation of postsynaptic opsins (spiking increase with excitatory and decrease with inhibitory opsins) was used as an internal reference for each electrode at each recording experiment. To compare neuron activity across treatments, we counted spikes for a 5 sec period before and after treatment. During this time period, spikes were counted when recorded extracellular signal surpassed nine standard deviations of baseline noise (ref). Data were not normally distributed, so non-parametric Kruskal-Wallis tests were used. Dunn’s post-hoc pairwise comparisons were performed only when global Kruskal-Wallis tests were significant, and no additional correction for electrode-level replication was applied as each electrode served as its own internal control via baseline and LED stimulation. Absolute *P* values are present in the manuscript and we considered *p* < 0.05 a useful threshold to reject the null hypothesis and 95% confidence level were used in each test. Statistical analyses were performed at the electrode level with multiple electrodes recorded per culture; electrodes from the same culture were treated as technical replicates.

For whole cell recordings, neurons with a resting potential above −40 mV and below −80 mV were excluded from analysis. Each N reflects a single recording of a single postsynaptic neuron. Each set of data collected and analyzed was assessed for normality using the Shapiro-Wilks test (p < 0.05). To compare three or more experimental groups we applied one-way ANOVA (parametric data, p < 0.05) with Tukey post-hoc analysis or Kruskal-Wallis testing (non-parametric data, p < 0.05) with Dunn’s post hoc. Comparisons of two experimental groups were assessed with either two-way Student’s t-tests (parametric, p < 0.05) or Welch’s unpaired t-test (non-parametric, p< 0.05). For comparisons of luciferase brightness in Persist-Int constructs, ANOVA testing with Fisher’s LSD post hoc was applied (p < 0.05). In Act-Int efficiency experiments, neurotransmitter and Act-Int data were compared with paired t-tests at each red light exposure time (p < 0.05, for parametric data: 4 ms, 16 ms, 32 ms) or a Wilcoxon Ranked Sums test (p < 0.05, nonparametric data: 2 ms). 95% confidence level were used in each test.

### Estimation of donor requirements for FRET-coupled Persist-Int

To assess whether short-range donor–opsin coupling could in principle support Persist-Int signaling, we calculated an optimistic lower-bound estimate for the number of active luciferase donor molecules required under a simplified FRET-only model. FRET efficiency was calculated from donor–acceptor separation *r* and Förster radius *R*_0_ as

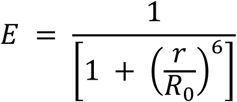

Using *R*_0_ = 5 nm and *r* = 5 nm to give *E* = 0.5. For each donor, the transferred excitation rate was taken as

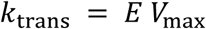

where *V*_max_ is donor photon output in photons s^−1^ molecule^−1^. To relate this to opsin activation, we used the characteristic deactivation time *τ*_off_ of the postsynaptic opsin as a simple integration window. Under this approximation, one donor contributes

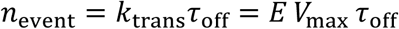

effective transferred excitations during one opsin open-state lifetime. The minimum number of concurrently active donor molecules required to reach one effective excitation per integration window is therefore

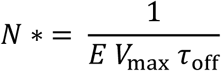

where *τ*_off_ is the opsin off-time in seconds. Values below 1 were reported as **<1**.

For example, for GeNL_eKL9h with *V*_max_ = 21.5 photons s^−1^ molecule^−1^ and ChR2(C128S) with *τ*_off_ = 106 s,

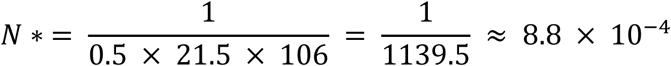

and this value was therefore reported as **<1**. This calculation was intended only to define an optimistic lower bound for donor requirements under favorable nanoscale coupling. It excludes radiative transfer and does not incorporate dipole orientation, spectral mismatch, dynamic geometry, or the true channel gating probability per transferred excitation.

## Supporting information

Supplementary text, tables, figures references, Peptide sequences

## Acknowledgements

We thank the members of the Bioluminescence Hub (http://www.bioluminescencehub.org/) for discussions and advice on experiments. This research was supported by grants from the National Science Foundation (NSF NeuroNex 1707352 to C.I.M., D.L., U.H., N.C.S.) and the US National Institutes of Health (R21MH101525 to U.H.; U01NS099709 to U.H., C.I.M., N.C.S.; R01NS120832 to U.H., C.I.M., N.C.S.; R21NS132089 to U.H.; R21NS135545 to U.H.; R21MH135326 to U.H.). Figures were generated using BioRender.com.

## Author Contributions

A.N.S., M.P., and R.S. designed and conducted experiments and analyzed resulting data. M.O.T. and G.G.L. performed experiments. E.L.C. and A.D.S. contributed to data analysis. N.C.S. performed theoretical modeling. N.C.S., C.I.M., and D.L. provided critical input throughout the studies. A.N.S. and U.H. wrote a draft of the manuscript. A.N.S., U.H., N.C.S. and D.L. generated the final manuscript. U.H. directed the overall study.

## Data and reagent availability

Raw and processed data sets from experiments performed in this study will be freely available via the Brown Digital Repository. Plasmids used in this study are being deposited for distribution by AddGene.

