## Supplementary text, tables, figures references, Peptide sequences for "Control of Synaptic Communication through Molecularly Engineered Bioluminescence Light Emission and Sensing"

### Supplement

#### Supplementary Text

#### Supplementary Tables

**Supplementary Table 1. Order-of-magnitude comparison of estimated luciferase-generated photon flux with literature opsin sensitivity benchmarks**
**Supplementary Table 2. Emitter flux divided by opsin benchmark flux for single-molecule isotropic emission at 20 nm distance**
**Supplementary Table 3. Plasmids used in these studies**
**Supplementary Table 4. Statistics**

#### Supplementary Figures

**S1. Whole cell, postsynaptic responses to light.**
**S2. Action potential characteristics across Act-Int and Persist-Int.**
**S3. Change in membrane voltage.**
**S4. Control whole cell recordings for split luciferase Act-Int.**
**S5. Potential influence of intravesicular pH on luciferase function in Act-Int experiments.**
**S6. Int synaptic transmission depends on synaptic connection.**

#### Supplementary References

#### Peptide Sequences

**Supplementary Text**

**Act-Int - Calculations of luciferase released from presynaptic neurons**

Parameters to be considered are the size/volume of the luciferase, the loading capacity of presynaptic vesicles, and the number of vesicles present in a mammalian synapse. The luciferase (hPOMC-sbGLuc-P2A) has a length of 217 amino acids, and thus a molecular weight of 23.19 kDa[1]. Assuming a theoretical spherical shape with a radius of R can encompass this peptide, we can use the molecular weight to calculate the volume the protein occupies[2], 27.9 $nm^3$.

We previously showed that hPOMC-sbGLuc is targeted to both synaptic vesicles (SVs) and large dense core vesicles (LDCVs)[3]. The diameter of synaptic vesicles is 40 nm which expands by 23.9% ± 4.8% when loaded with neurotransmitters to a size of 47.64 – 51.48 nm[4]. The diameter of LDCVs is between 111.5 - 195.9 nm[5]. Accordingly, the total volume available for cargo is 5.66 – 7.14 x $10^4$ $nm^3$ for SVs and 72.6 – 393.6 x $10^4$ $nm^3$ for LDCVs.

In addition to the volume capacity of the individual vesicle, the final volume available to hold luciferases is determined by the number of vesicles present in a synapse. In the mammalian synapse about 200 – 500 SVs are present in the terminal with 10 – 30 in the active zone[6]. This provides a holding volume for luciferases of 113 – 357 x $10^5$ $nm^3$ in the terminal and 5.6 – 21.4 x $10^5$ $nm^3$ in the active zone. With LDCVs comprising only 1 – 2 % of the number of SVs in most locations in the brain[5], 2 – 10 can be found in the terminal and 0.1 – 0.6 (rounded up to 1) in the active zone. Accordingly, the total holding capacity for luciferases in LDCVs 14.5 – 394 x $10^5$ $nm^3$ in the terminal and 7.2 – 39.3 x $10^5$ $nm^3$ in the active zone.

With the calculated volumes occupied by the luciferase and provided by the vesicles together with the number of vesicles present at the terminal and the active zone, the total number of luciferase molecules available in a synapse can be estimated at, for SVs, 41 – 130 x $10^4$ (terminal) and 2 – 7.7 x $10^4$ (active zone), and for LDCVs, 5.2 – 140 x $10^4$ (terminal) and 2.6 – 14 x $10^4$ (active zone). Based on this, in a single synapse a maximum combined 5 x $10^5$-16 x $10^5$ molecules of sbGLuc are released in a single one-second red light stimulation of the presynaptic neuron. These values assume that a one second stimulation induces the release of all available vesicles from the presynaptic neuron and that each vesicle is filled completely with luciferase molecules. In reality, loading volume will be diminished due to co-localization with innate neurotransmitters, the aqueous environment in the vesicle and ions taking up space as well as possibly causing steric hindrances. This compares to NT release with 8000 glutamate molecules in one synaptic vesicle[7]: 160 x$10^4$-400x$10^4$ (reserve pool) and 8x$10^4$-24x$10^4$ (active zone), combined 16x$10^5$-42x$10^5$ molecules of glutamate being released in a single one-second red light stimulation of the presynaptic neuron.

#### Supplementary Tables

##### Supplementary Table 1. Order-of-magnitude comparison of estimated luciferase-generated photon flux with literature opsin sensitivity benchmarks

| Class | Molecule / benchmark | λ (nm) | Input value | Flux at 5 nm (ph $s^{-1}$ $mm^{-2}$) | Flux at 10 nm (ph $s^{-1}$ $mm^{-2}$) | Flux at 20 nm (ph $s^{-1}$ $mm^{-2}$) | Notes |
|---|---|---|---|---|---|---|---|
| Emitter | NanoLuc + furimazine | 460 | 8.6 photons $s^{-1}$ $molecule^{-1}$ | $2.74 \times 10^{10}$ | $6.84 \times 10^{9}$ | $1.71 \times 10^{9}$ | Absolute per-molecule output from calibrated purified NanoLuc. |
| Emitter | GeNL_eKL9h + furimazine | 460 | 21.5 photons $s^{-1}$ $molecule^{-1}$ | $6.84 \times 10^{10}$ | $1.71 \times 10^{10}$ | $4.28 \times 10^{9}$ | Moderately constrained inference from the directed-evolution path placing eKL9h below SSLuc. |
| Emitter | SSLuc / GeNL-SSLuc + furimazine | 460 / 520 | 86 photons $s^{-1}$ $molecule^{-1}$ | $2.74 \times 10^{11}$ | $6.84 \times 10^{10}$ | $1.71 \times 10^{10}$ | Anchored to NanoLuc absolute calibration and 10 ± 1-fold higher in vitro steady-state output with furimazine. |
| Opsin benchmark | ChR2(C128S) | 470 | 0.01 mW $mm^{-2}$ | - | - | $2.37 \times 10^{13}$ | Highly light-sensitive step-function channelrhodopsin benchmark. |
| Opsin benchmark | ChRger3 | ~500 | 0.008 mW $mm^{-2}$ | - | - | $2.01 \times 10^{13}$ | Low-light benchmark from source assay; not a universal ED50. |
| Opsin benchmark | CheRiff | 460 | 0.22 mW $mm^{-2}$ | - | - | $5.09 \times 10^{14}$ | Converted from 22 ± 10 mW $cm^{-2}$ benchmark in source assay. |
| Opsin benchmark | ChRmine | 585 | 0.08 mW $mm^{-2}$ | - | - | $2.36 \times 10^{14}$ | Source-assay spiking benchmark; not a universal ED50. |
| Opsin benchmark | hGtACR2 | 470 | 0.05 mW $mm^{-2}$ | - | - | $1.18 \times 10^{14}$ | EPD50 benchmark reported in later optophysiology literature. |
| Opsin benchmark | ChrimsonR | 590 | 1.0 mW $mm^{-2}$ | - | - | $2.97 \times 10^{15}$ | Conservative red-light operational benchmark. |

Emitter flux[8] was estimated by treating each luciferase as an isotropic point source and applying the inverse-square law, $\Phi(r) = N\gamma/(4\pi r^2)$, where Nγ is the photon emission rate in photons $s^{-1}$ $molecule^{-1}$ and r is distance. Opsin-side values were converted from literature irradiance benchmarks[9–16] to photon flux using $\Phi = P\lambda/(hc)$. Because reported opsin sensitivities depend on assay endpoint, wavelength, pulse duration, expression level, and cell type, these values are presented as order-of-magnitude operational benchmarks rather than standardized cross-study ED50 constants.

**Supplementary Table 2. Emitter flux divided by opsin benchmark flux for single-molecule isotropic emission at 20 nm distance**

| Emitter estimate | ChR2(C128S) | ChRger3 | CheRiff | ChRmine | hGtACR2 | ChrimsonR |
|---|---|---|---|---|---|---|
| NanoLuc at 20 nm | $7.23 \times 10^{-5}$ | $8.50 \times 10^{-5}$ | $3.36 \times 10^{-6}$ | $7.26 \times 10^{-6}$ | $1.45 \times 10^{-5}$ | $5.76 \times 10^{-7}$ |
| GeNL_eKL9h at 20 nm | $1.81 \times 10^{-4}$ | $2.12 \times 10^{-4}$ | $8.40 \times 10^{-6}$ | $1.82 \times 10^{-5}$ | $3.62 \times 10^{-5}$ | $1.44 \times 10^{-6}$ |
| SSLuc / GeNL-SSLuc at 20 nm[a] | $7.23 \times 10^{-4}$ | $8.50 \times 10^{-4}$ | $3.36 \times 10^{-5}$ | $7.26 \times 10^{-5}$ | $1.45 \times 10^{-4}$ | $5.76 \times 10^{-6}$ |

Values are reported as fold shortfall (Φ_opsin / Φ_emitter) at 20 nm assuming isotropic emission. Values are rounded to 1–2 significant figures. Across all emitter–opsin pairs, isotropic single-molecule emission falls several orders of magnitude below literature activation benchmarks. These estimates represent a lower-bound radiative comparison and do not account for nanoscale proximity, local clustering, prolonged integration by highly light-sensitive opsins, or more efficient near-field transfer mechanisms than simple isotropic radiation (i.e., FRET).

**Supplementary Table 3. Plasmids used in these studies**

| Plasmid | Figure | Addgene # |
|---|---|---|
| Presynaptic light emitters | | |
| Act-Int | | |
| pcDNA CAG POMC sbGLuc P2A Chrimson dTom | 3 E-H<br>5 B, D, E<br>6<br>7 E, G-I | 256449 |
| pcDNA CAG Synb sbGLuc P2A ChrimsonR dTom | 5 B | 256450 |
| pcDNA CAG VGAT sbGluc P2A ChrimsonR dTom | 5 B | 256451 |
| pcDNA CAG POMC mCherry 22.0_RLuc8.6 | 5 D | 256452 |
| pcDNA CAG POMC GeNL_SSLuc P2A ChrimsonR dTom | 5 D | 256453 |
| pcDNA CAG POMC ssLuc P2A ChrimsonR dTom | 5 D | 256454 |
| pcDNA CAG POMC linker SSLuc_smb P2A ChrimsonR dTom | 5 D | 256455 |
| Persist-Int | | |
| pcDNA CAG GeNL_eKL9h linker ICAM Nrxn3b P2A dTom | 2<br>3 C-G<br>4 B, E<br>7 K | 256456 |
| pcDNA CAG GeNL_SS linker ICAM Nrxn3b P2A dTom | 3 H<br>4 B, D, F<br>7 K | 256457 |
| pcDNA CAG GeNL_SS linker CD4 Nrxn3b P2A dTom | 4 D<br>7 K | 256458 |
| pcDNA CAG NanoLuc-ICAM-Nrxn3b | 4 B | 256459 |
| pcDNA CMV ssLuc linker B7 EYFP | 4 D | 256460 |
| Presynaptic opsins (co-expressed with light emitter) | | |
| pcDNA CAG ChrimsonR dTom | 4 B, D-F<br>7 K | 256461 |
| pcDNA CAG ChrimsonR EYFP | 4 D (SSLuc-B7) | 256462 |
| pcDNA CAG ChR2(C128S) dTom | 5 D (mCh-RLuc8.6) | 256463 |
| Postsynaptic opsins | | |
| pcDNA CAG hGtACR2 EYFP | 2 | 256465 |
| pcDNA CAG ChR2(C128S) EYFP | 2<br>3 E-H<br>4 B, D-F<br>5 B, D, E<br>6<br>7 E, K | 256466 |
| pcDNA CAG Cheriff EYFP | 3 C-G<br>4 E<br>5 E | 256467 |
| pcDNA CAG ChRmine EYFP | 3 E-F<br>4 E<br>7 K | 256468 |
| pcDNA CAG SP VChR1 ts-EYFP-er | 3 E-F<br>4 E | 256469 |

| | | |
|---|---|---|
| pcDNA CAG ChrimsonR dTom | 4 F | 256461 |
| pcDNA CAG ChR2(C128S) dTom | 4 D (SSLuc-B7) | 256463 |
| | 7 H-I | |
| pcDNA CAG ChrimsonR EYFP | 5 D | 256462 |
| | 7 K | |
| pcDNA CAG SSLuc_lgb GGS 15 aa linker ChRmine EYFP | 5 D | 256470 |
| pcDNA CMV ChRger3 TS EYFP | 5 E | 256471 |
| $Ca^{2+}$ sensor | | |
| pcDNA CAG jGCaMP8s | 7 H-I | 256472 |

#### Supplementary Table 4. Statistics

| Figure | Panel | Group<br>(Group #: N, Mean, SEM) | Shapiro-Wilk | Statistical Test | Overall Values | Group Comparisons | Post Hoc<br>($\alpha$ < 0.05) |
|---|---|---|---|---|---|---|---|
| 2 | C | MEA GeNL_eKL9h-ICAM x ChR2(CS) LED<br>(1: 62, 7.039, 1.002) | W = 0.6842<br>P < 0.0001 | Kruskal-Wallis + Dunn's | H = 101.0<br>P < 0.0001 | 1:3 | Mean Rank: 71.29<br>P < 0.0001 |
| | | MEA GeNL_eKL9h-ICAM x ChR2(CS) Luciferin (2: 16, 1.598, 0.0721) | W = 0.8643<br>p = 0.0222 | | | 1:2 | Mean Rank: 28.23<br>p = 0.024 |
| | | MEA GeNL_eKL9h-ICAM x ChR2(CS) vehicle<br>(3: 53, 0.7199, 0.01995) | W = 0.9890<br>p = 0.9057 | | | 2:3 | Mean Rank: 43.06<br>p = 0.0002 |
| | | MEA GeNL_eKL9h-ICAM x hGtACR2 LED<br>(1: 38, 0.3813, 0.03651) | W = 0.9660<br>p = 0.2952 | Kruskal-Wallis + Dunn's | H = 78.05<br>P < 0.0001 | 1:3 | Mean Rank: -26.59<br>p = 0.0013 |
| | | MEA GeNL_eKL9h-ICAM x hGtACR2 Luciferin (2: 38, 0.02291, 0.01051) | W = 0.4063<br>P < 0.0001 | | | 1:2 | Mean Rank: 39.58<br>P < 0.0001 |
| | | MEA GeNL_eKL9h-ICAM x hGtACR2 vehicle<br>(3: 38, 0.6710, 0.03545) | W = 0.9650<br>p = 0.2756 | | | 2:3 | Mean Rank: -66.17<br>P < 0.0001 |
| 3 | E | NT_Half-width<br>(1: 43, 0.5266, 0.02888) | W = 0.7393<br>P < 0.0001 | Wilcoxon Signed Rank (two tailed) | W = 578<br>p = 0.0003 | | |
| | | Int_Half-width<br>(2: 43, 0.5588, 0.02372) | W = 0.9158<br>p = 0.0039 | | | | |
| | | NT_AHP<br>(1: 43, -5.901, 0.6198) | W = 0.9279<br>p = 0.0098 | Wilcoxon Signed Rank (two tailed) | W = 938<br>P < 0.0001 | | |
| | | Int_AHP<br>(2: 43, 1.345, 0.4747) | W = 0.8006<br>P < 0.0001 | | | | |
| | G | Act-Int (POMC sbGLuc) x ChR2(CS)<br>(1: 5, 85.04, 34.66) | W = 0.8459<br>p = 0.1818 | ANOVA + Tukey's | F = 1.374<br>p = 0.2864 | 1:2 | p = 0.9718 |
| | | | | | | 1:3 | p = 0.7980 |
| | | Act-Int (POMC sbGLuc) x CheRiff<br>(2: 5, 102.7, 28.48) | W = 0.8467<br>p = 0.1844 | | | 1:4 | p = 0.7339 |
| | | Persist-Int (GeNL_eKL9h-ICAM) x ChR2(CS)<br>(3: 5, 122.2, 32.63) | W = 0.8394<br>p = 0.1634 | | | 2:3 | p = 0.9624 |
| | | | | | | 2:4 | p = 0.4799 |
| | | Persist-Int (GeNL_eKL9h-ICAM) x CheRiff<br>(4: 5, 43.10, 15.11) | W = 0.9114<br>p = 0.4762 | | | 3:4 | p = 0.2492 |
| | H | Act-Int (POMC sbGLuc x ChR2(CS))<br>(1: 13, 26.63, 2.891) | W = 0.9417<br>p = 0.4789 | Unpaired Welch's t-test (two tailed) | t = 0.06895<br>df = 6.761<br>p = 0.9470 | | |
| | | Persist-Int (GeNL_SS-ICAM x ChR2(CS))<br>(2: 5, 26.22, 5.091) | W = 0.9247<br>p = 0.5608 | | | | |
| 4 | B | GeNL_SS-ICAM x ChR2(CS)<br>(1: 5, 26.22, 5.091) | Above<br>(4H.2) | ANOVA + Fisher's LSD | F = 11.38<br>p = 0.0014 | 1:2 | p = 0.0369 |
| | | GeNL_eKL9h-ICAM x ChR2(CS)<br>(2: 6, 15.10, 3.003) | W = 0.9482<br>p = 0.7254 | | | 1:3 | p = 0.0004 |
| | | NanoLuc-ICAM x ChR2(CS)<br>(3: 5, 2.392, 1.075) | W = 0.8668<br>p = 0.2536 | | | 2:3 | p = 0.0198 |
| | D | GeNL_SS-ICAM x ChR2(CS)<br>(1: 5, 26.22, 5.091) | Above<br>(4H.2) | ANOVA + Tukey's | F = 17.82 | 1:2 | p = 0.0037 |

| | | | | | | | |
|---|---|---|---|---|---|---|---|
| | | GeNL_SS-CD4 x ChR2(CS) (2: 5, 8.172, 1.681) | W = 0.9752 p = 0.9074 | | p = 0.0003 | 1:3 | p = 0.0002 |
| | | SSLuc-B7 x ChR2(CS) (3: 5, 0.8080, 0.2049) | W = 0.8853 p = 0.3338 | | | 2:3 | p = 0.2520 |
| | E | GeNL_eKL9h-ICAM x ChR2(CS) (1: 6, 15.10, 3.003) | Above (5D.2) | ANOVA + Tukey's | F = 0.4908 p = 0.6933 | 1:2 | p = 0.9721 |
| | | | | | | 1:3 | p = 0.9496 |
| | | GeNL_eKL9h-ICAM x ChRmine (2: 5, 13.11, 2.710) | W = 0.9397 p = 0.6640 | | | 1:4 | p = 0.8945 |
| | | GeNL_eKL9h-ICAM x VChR1 (3: 6, 17.44, 3.866) | W = 0.9272 p = 0.5589 | | | 2:3 | p = 0.7834 |
| | | | | | | 2:4 | p = 0.7136 |
| | | GeNL_eKL9h-ICAM x CheRiff (4: 4, 18.54, 3.068) | W = 0.8912 p = 0.3889 | | | 3:4 | p = 0.9958 |
| | F | GeNL_SS-ICAM x ChR2(CS) (1: 5, 26.22, 5.091) | Above (4H.2) | Unpaired Welch's t-test (two tailed) | t = 2.445 df = 7.632 p = 0.0416 | | |
| | | GeNL_SS-ICAM x ChrimsonR (2: 5, 42.17, 4.073) | W = 0.9247 p = 0.9642 | | | | |
| 5 | B | POMC sbGLuc x ChR2(CS) (1: 13, 26.63, 2.891) | Above (4H.1) | Kruskal-Wallis + Dunn's | H = 1.531 p = 0.4651 | 1:2 | P > 0.9999 |
| | | Synb-sbGLuc x ChR2(CS) (2: 5, 21.59, 3.756) | W = 0.9183 p = 0.5190 | | | 1:3 | P > 0.9999 |
| | | VGAT-sbGLuc x ChR2(CS) (3: 5, 27.92, 2.301) | W = 0.7637 p = 0.0397 | | | 2:3 | p = 0.6762 |
| | D | POMC sbGLuc x ChR2(CS) (1: 13, 26.63, 2.891) | Above (4H.1) | Kruskal-Wallis + Dunn's | H = 14.09 p = 0.0070 | 1:2 | p = 0.8383 |
| | | | | | | 1:3 | P > 0.9999 |
| | | POMC mCh22.0_RLuc8.6 x ChrimsonR (2: 5, 14.18, 4.075) | W = 0.9383 p = 0.6540 | | | 1:4 | p = 0.0545 |
| | | | | | | 1:5 | P > 0.9999 |
| | | POMC GeNL_SS x ChR2(CS) (3: 7, 15.18, 6.070) | W = 0.8834 p = 0.2420 | | | 2:3 | P > 0.9999 |
| | | | | | | 2:4 | P > 0.9999 |
| | | POMC SSLuc x ChR2(CS) (4: 6, 5.732, 5.404) | W = 0.5902 P < 0.0001 | | | 2:5 | p = 0.2361 |
| | | | | | | 3:4 | P > 0.9999 |
| | | POMC SSLuc$_{smb}$ x SSLuc$_{lgb}$-ChRmine (5: 6, 31.76, 4.184) | W = 0.9488 p = 0.7309 | | | 3:5 | p = 0.3826 |
| | | | | | | 4:5 | p = 0.0150 |
| | E | POMC sbGLuc x ChR2(CS) (1: 13, 26.63, 2.891) | Above (4H.1) | ANOVA + Tukey's | F = 0.2350 p = 0.7928 | 1:2 | p = 0.7752 |
| | | POMC sbGLuc x ChRger3 (2: 5, 23.08, 4.844) | W = 0.9620 p = 0.8217 | | | 1:3 | p = 0.9873 |
| | | POMC sbGLuc x CheRiff (3: 5, 25.84, 2.895) | W = 0.9564 p = 0.7829 | | | 2:3 | p = 0.8982 |
| 6 | C | NT_2ms (1: 5, 1.500, 0.4000) | W = 0.7709 p = 0.0460 | Wilcoxon Matched-Pairs Sign | W = -5.000 | | |

| | | | | | | | |
|---|---|---|---|---|---|---|---|
| | | Int_2ms<br>(2: 5, 1.200, 0.5831) | W = 0.9020<br>p = 0.4211 | Rank (two way) | p = 0.7500 | | |
| | | NT_4ms<br>(3: 3, 1.000, 0.5774) | W = 1.000<br>P > 0.9999 | Paired t-test (two way) | t = 0.5000<br>df = 2<br>p = 0.6667 | | |
| | | Int _4ms<br>(4: 3, 1.667, 0.8819) | W = 0.9643<br>p = 0.6369 | | | | |
| | | NT_16ms<br>(5: 3, 5.667, 2.603) | W = 0.9959<br>p = 0.8776 | Paired t-test (two way) | t = 0.3273<br>df = 2<br>p = 0.7745 | | |
| | | Int _16ms<br>(6: 3, 4.667, 2.186) | W = 0.8547<br>p = 0.2530 | | | | |
| | | NT_32ms<br>(7: 4, 27.00, 12.73) | W = 0.8879<br>p = 0.3737 | Paired t-test (two way) | t = 0.02020<br>df = 3<br>p = 0.9851 | | |
| | | Int _32ms<br>(8: 4, 26.50, 12.82) | W = 0.9265<br>p = 0.5740 | | | | |
| 7 | E | connected*<br>(1: 15, 81.45, 14.28) | W = 0.9065<br>p = 0.1198 | Unpaired Welch's t-test (two way) | t = 5.463<br>df = 14.2<br>P < 0.0001 | | |
| | | nearby*<br>(2: 5, 3.091, 1.375) | W = 0.8927<br>p = 0.3709 | | | | |
| | H | jGCaMP8s LED before<br>(1: 12, 0.3975, 0.1385) | W = 0.7948<br>p = 0.0082 | Kruskal-Wallis + Dunn's | W = 26.54<br>P < 0.0001 | 1:2 | P < 0.0001 |
| | | jGCaMP8s LED after<br>(2: 6, 10.81, 0.7178) | W = 0.9692<br>p = 0.8871 | | | 1:3 | p = 0.2438 |
| | I | jGCaMP8s CTZ before<br>(1: 12, 0.3975, 0.1385) | Above<br>(8H.1) | | | 2:3 | p = 0.1085 |
| | | jGCaMP8s CTZ after<br>(3: 6, 2.385, 0.1520) | W = 0.9500<br>p = 0.7404 | | | | |
| | K | connected | W = 0.9233<br>p = 0.1147 | Mann-Whitney | U = 0<br>p < 0.0001 | | |
| | | nearby | W = 0.7331<br>p = 0.0053 | | | | |
| S1 | A | GeNL_eKL9h-ICAM x CheRiff blue light<br>(1: 4, 11.80, 3.386) | W = 0.9806<br>p = 0.9058 | Kruskal-Wallis + Dunn's | W = 34.40<br>p = 0.0113 | 2:8 | p = 0.0722 |
| | | GeNL_eKL9h-ICAM x ChR2(CS) blue light<br>(2: 6, 6.160, 1.968) | W = 0.7513<br>p = 0.0205 | | | 2:14 | p = 0.6426 |
| | | GeNL_eKL9h-ICAM x ChRmine blue light<br>(3: 5, 8.590, 1,810) | W = 0.9264<br>p = 0.5717 | | | 8:9 | p = 0.0735 |
| | | GeNL_eKL9h-ICAM x VChR1 blue light<br>(4: 6, 9.098, 1.877) | W = 0.9619<br>p = 0.8346 | | | 8:10 | p = 0.4373 |
| | | GeNL_SS-ICAM x ChR2(CS) blue light<br>(5: 5, 8.478, 1.476) | W = 0.9864<br>p = 0.9656 | | | 8:11 | p = 0.0779 |
| | | GeNL_SS-CD4 x ChR2(CS) blue light<br>(6: 5, 8.764, 2.324) | W = 0.9031<br>p = 0.4274 | | | 8:12 | p = 0.0839 |

| | | | | | | | |
|---|---|---|---|---|---|---|---|
| | | SSLuc-B7 x ChR2(CS) blue light (7: 5, 7.252, 1.896) | W = 0.9669 p = 0.8549 | | | 8:13 | p = 0.2448 |
| | | GeNL_SS-ICAM x ChrimsonR blue light (8: 5, 20.94, 2.248) | W = 0.9564 p = 0.7825 | | | 8:16 | p = 0.7637 |
| | | NanoLuc-ICAM x ChR2(CS) blue light (9: 5, 4.934, 1.221) | W = 0.9134 p = 0.4885 | | | 8:17 | p = 0.3731 |
| | | Synb-sbGLuc x ChR2(CS) blue light (10: 5, 6.136, 1.386) | W = 0.9501 p = 0.7379 | | | 8:18 | p = 0.2025 |
| | | VGAT-sbGLuc x ChR2(CS) blue light (11: 5, 5.134, 1.086) | W = 0.8729 p = 0.2785 | | | 9:14 | p = 0.5993 |
| | | POMC SSLuc x ChR2(CS) blue light (12: 6, 5.318, 1.163) | W = 0.7137 p = 0.0086 | | | 11:14 | p = 0.6293 |
| | | POMC GeNL_SS x ChR2(CS) blue light (13: 7, 6.666, 1.649) | W = 0.8534 p = 0.1320 | | | 12:14 | p = 0.7293 |
| | | POMC mCh22.0_RLuc8.6 x ChrimsonR blue light (14: 5, 14.80, 1.501) | W = 0.8334 p = 0.1476 | | | All other comparisons | P > 0.9999 |
| | | POMC sbGLuc x Rh2 blue light (15: 5, 7.636, 0.9707) | W = 0.9237 p = 0.5540 | | | | |
| | | POMC sbGLuc x ChRger3 blue light (16: 5, 6.660, 2.303) | W = 0.6361 p = 0.0018 | | | | |
| | | POMC sbGLuc x ChR2(CS) blue light (17: 13, 8.008, 1.338) | W = 0.8497 p = 0.0282 | | | | |
| | | POMC sbGLuc x CheRiff blue light (18: 4, 5.830, 2.301) | W = 0.8596 p = 0.2588 | | | | |
| | | POMC SSLuc$_{smb}$ x ChRmine blue light (19: 6, 10.50, 1.983) | W = 0.8135 p = 0.0775 | | | | |
| | B | GeNL_eKL9h-ICAM x CheRiff membrane potential (1: 4, -54.70, 1.501) | W = 0.9497 p = 0.7143 | Kruskal-Wallis + Dunn's | W = 17.59 p = 0.4832 | 12:14 | p = 0.8567 |
| | | GeNL_eKL9h-ICAM x ChR2(CS) membrane potential (2: 6, -51.19, 2.114) | W = 0.9220 p = 0.5196 | | | All other comparisons | p > 0.9999 |
| | | GeNL_eKL9h-ICAM x ChRmine membrane potential (3: 5, -50.32, 3.119) | W = 0.9890 p = 0.9763 | | | | |
| | | GeNL_eKL9h-ICAM x VChR1 membrane potential (4: 6, -52.95, 1.639) | W = 0.9458 p = 0.7064 | | | | |
| | | GeNL_SS-ICAM x ChR2(CS) membrane potential (5: 5, -52.67, 3.598) | W = 0.9783 p = 0.9252 | | | | |
| | | GeNL_SS-CD4 x ChR2(CS) membrane potential (6: 5, -52.34, 4.509) | W = 0.8119 p = 0.1010 | | | | |
| | | SSLuc-B7 x ChR2(CS) membrane potential (7: 5, -57.57, 2.680) | W = 0.9082 p = 0.4570 | | | | |
| | | GeNL_SS-ICAM x ChrimsonR membrane potential (8: 5, -55.79, 5.203) | W = 0.8927 p = 0.3707 | | | | |
| | | NanoLuc-ICAM x ChR2(CS) membrane potential (9: 5, -61.35, 4.693) | W = 0.9193 p = 0.5252 | | | | |
| | | Synb-sbGLuc x ChR2(CS) membrane potential (10: 5, -57.36, 4.817) | W = 0.9236 p = 0.5532 | | | | |
| | | VGAT-sbGLuc x ChR2(CS) membrane potential (11: 5, -50.33, 2.215) | W = 0.9294 p = 0.5921 | | | | |
| | | POMC SSLuc x ChR2(CS) membrane potential (12: 6, -63.01, 4.517) | W = 0.9917 p = 0.9929 | | | | |

| | | | | | | | |
|---|---|---|---|---|---|---|---|
| | | POMC GeNL_SS x ChR2(CS) membrane potential (13: 7, -53.73, 4.010) | W = 0.9064<br>p = 0.3716 | | | | |
| | | POMC mCh22.0_RLuc8.6 x ChrimsonR membrane potential (14: 5, -47.87, 1.786) | W = 0.9665<br>p = 0.8526 | | | | |
| | | POMC sbGLuc x Rh2 membrane potential (15: 5, -55.07, 3.715) | W = 0.9025<br>p = 0.4241 | | | | |
| | | POMC sbGLuc x ChRger3 membrane potential (16: 5, -57.83, 3.730) | W = 0.9593<br>p = 0.8028 | | | | |
| | | POMC sbGLuc x ChR2(CS) membrane potential (17: 13, -53.78, 2.245) | W = 0.8899<br>p = 0.0973 | | | | |
| | | POMC sbGLuc x CheRiff membrane potential (18: 5, -56.22, 3.949) | W = 0.9615<br>p = 0.8183 | | | | |
| | | POMC $SSLuc_{smb}$ x ChRmine membrane potential (19: 5, -54.51, 3.004) | W = 0.7577<br>p = 0.0350 | | | | |
| S2 | A | POMC sbGLuc x ChR2(CS) half-width (1: 7, 0.6106, 0.0611) | W = 0.7867<br>p = 0.0302 | Kruskal-Wallis + Dunn's | W = 2.790<br><br>p = 0.7323 | All comparisons | p > 0.9999 |
| | | POMC sbGLuc x CheRiff half-width (2: 7, 0.5787, 0.05290) | W = 0.8289<br>p = 0.0781 | | | | |
| | | GeNL_eKL9h-ICAM x ChR2(CS) half-width (3: 7, 0.5031, 0.06887) | W = 0.8370<br>p = 0.0931 | | | | |
| | | GeNL_eKL9h-ICAM x CheRiff half-width (4: 7, 0.5851, 0.07672) | W = 0.8880<br>p = 0.2643 | | | | |
| | | GeNL_eKL9h-ICAM x ChRmine half-width (5: 7, 0.4903, 0.03074) | W = 0.9394<br>p = 0.6333 | | | | |
| | | GeNL_eKL9h-ICAM x VChR1 half-width (6: 8, 0.575, 0.05634) | W = 0.8433<br>p = 0.0814 | | | | |
| | B | POMC sbGLuc x ChR2(CS) AHP (1: 7, 1.774, 1.035) | W = 0.8826<br>p = 0.2383 | Kruskal-Wallis + Dunn's | W = 3.820<br><br>p = 0.5757 | All comparisons | p > 0.9999 |
| | | POMC sbGLuc x CheRiff AHP (2: 7, 2.903, 1.984) | W = 0.7620<br>p = 0.0169 | | | | |
| | | GeNL_eKL9h-ICAM x ChR2(CS) AHP (3: 7, 0.5609, 1,060) | W = 0.9068<br>p = 0.3744 | | | | |
| | | GeNL_eKL9h-ICAM x CheRiff AHP (4: 7, 0.7339, 0.6964) | W = 0.7828<br>p = 0.0276 | | | | |
| | | GeNL_eKL9h-ICAM x ChRmine AHP (5: 7, -0.07289, 0.3344) | W = 0.9432<br>p = 0.6680 | | | | |
| | | GeNL_eKL9h-ICAM x VChR1 AHP (6: 8, 2.068, 1.206) | W = 0.8840<br>p = 0.2056 | | | | |
| S4 | | POMC sbGLuc x ChR2(CS) (1: 13, 26.63, 2.891) | Above (4H.1) | ANOVA + Tukey's | F = 14.46<br><br>p = 0.0001 | 1:2 | p = 0.5326 |
| | | POMC $SSLuc_{smb}$ x $SSLuc_{lgb}$-ChRmine (2: 6, 31.76, 4.184) | Above (6D.5) | | | 1:3 | p = 0.0003 |
| | | POMC $SSLuc_{smb}$ x $SSLuc_{lgb}$-ChRmine (no red light) (3; 4, 0.5425. 0.1545) | W = 0.9005<br>p = 0.4337 | | | 2:3 | p = 0.0002 |

#### Supplementary Figures

#### Supplementary Figure S1

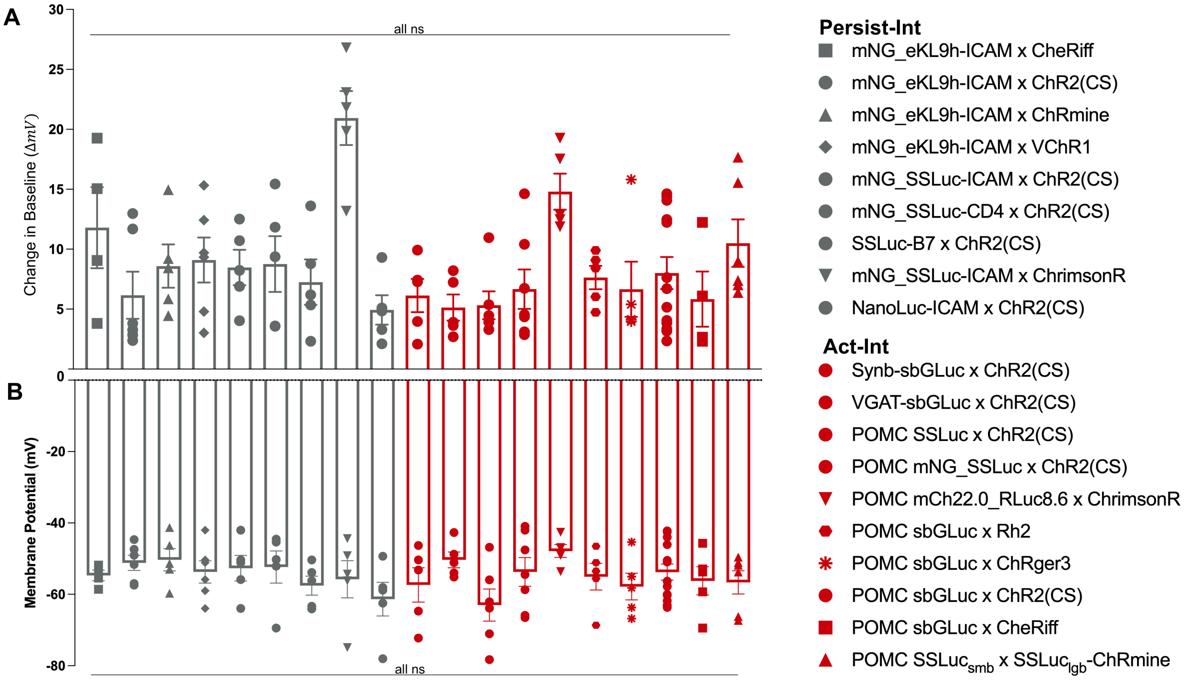

**Fig. S1. Whole cell, postsynaptic responses to light** (metal halide light source, blue filter). For every postsynaptic patch clamp recording used in analysis, the standard optogenetic response to light and the membrane potential was measured. **A** The change in baseline membrane potential (ΔmV) induced by direct light stimulation was not significantly different across postsynaptic recordings regardless of opsin type or Interluminescence type. **B** Membrane potential (mV) was not significantly different across all postsynaptic recordings regardless of light receptor type or Interluminescence paradigm.

#### Supplementary Figure S2

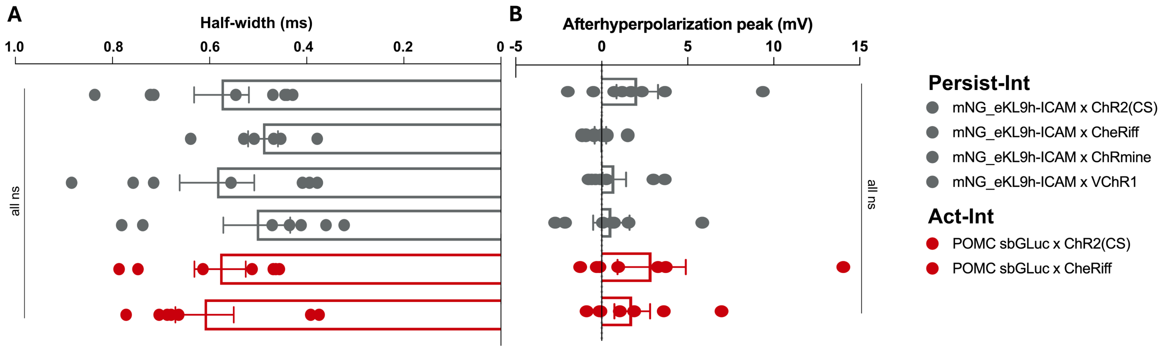

**Fig. S2. Action potential characteristics across Act-Int and Persist-Int.** Half-width (ms) and afterhyperpolarization peak (AHP, mV) were measured for the first action potential elicited with Int of various Act-Int and Persist-Int constructs. **A** Half-width was not significantly different for any construct tested regardless of Interluminescence type. **B** AHP was not significantly different for any construct tested regardless of Interluminescence type.

#### Supplementary Figure S3

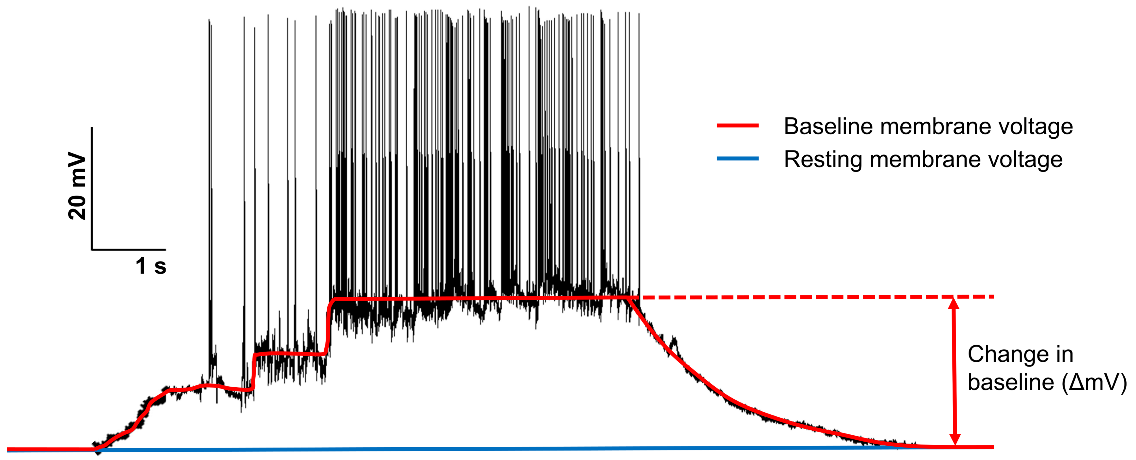

**Fig. S3. Change in membrane voltage.** To compare the efficacy of the various constructs tested we quantified the change in base membrane potential induced by each method. The change in baseline induced by stimulation (ΔmV) was defined by the difference between resting membrane potential (i.e., membrane potential before stimulus onset) and shifted baseline membrane potential at the saturation level during stimulation.

#### Supplementary Figure S4

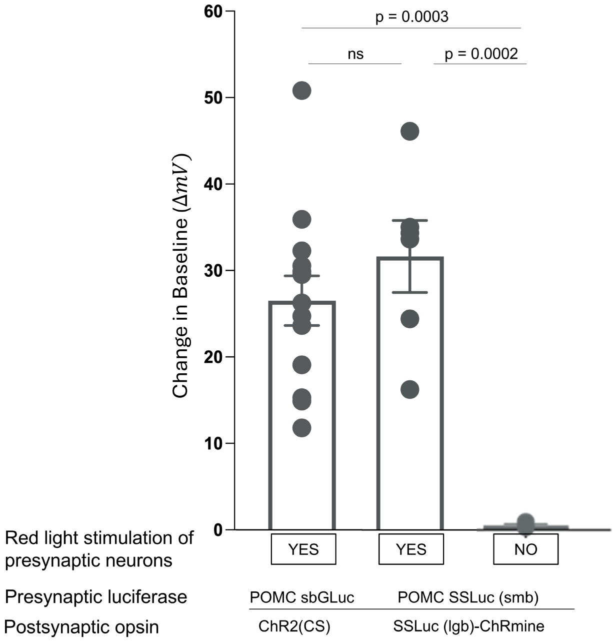

**Fig. S4. Control whole cell recordings for split luciferase Act-Int.** Whole cell recordings from co-cultures of pre- and postsynaptic neurons in the presence of luciferin and synaptic blockers and with (POMC sbGLuc (pre)/ChR2(C128S) (post), POMC $SSLuc_{smb}$ (pre)/$SSLuc_{lgb}$-ChRmine (post)) or without (POMC $SSLuc_{smb}$ (pre)/$SSLuc_{lgb}$-ChRmine (post)) red light stimulation of presynaptic neurons. No change in baseline membrane voltage (ΔmV) was measured in co-cultures when presynaptic neurons were not active (no red light stimulation), indicating that the small bit of the split luciferase, $SSLuc_{smb}$, requires vesicle fusion and release from activated neurons. N = 13 recordings (sbGLuc), 6 recordings ($SSLuc_{smb}$), 4 recordings ($SSLuc_{smb}$ w/o red light); Kruskal-Wallis; sbGLuc vs $SSLuc_{smb}$ $p = 0.5326$; sbGLuc vs $SSLuc_{smb}$ w/o red light $p = 0.0003$; $SSLuc_{smb}$ vs $SSLuc_{smb}$ w/o red light $p = 0.0002$.

#### Supplementary Figure S5

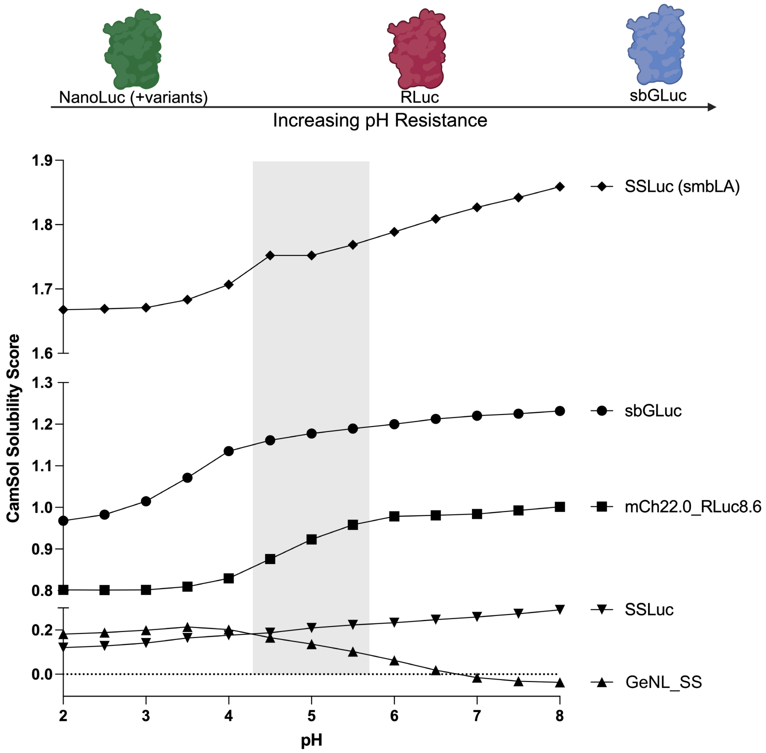

**Fig. S5. Potential influence of intravesicular pH on luciferase function in Act-Int experiments**. To quantitatively model pH sensitivities of the various luciferases tested in Act-Int (sbGLuc, mCh22.0_RLuc8.6, GeNL_SS, SSLuc, and $SSLuc_{smb}$) we ran luciferase sequences (with POMC on the N-terminus and the 5' end of P2A after cleavage included to the C-terminus where applicable) through CamSol software[17,18] to yield peptide solubility scores for pH 2 to 8. Lower scores indicate lower solubility at a given pH, which can indicate that a protein will misfold and aggregate, making it nonfunctional. As expected, sbGLuc and mCh22.0_RLuc8.6 showed higher solubility compared to NanoLuc variants SSLuc and GeNL_SS at vesicular pHs, concurrent with their respective pH resistances (top). The small bit of SSLuc had the highest solubility score at pH 5, ~1.5x that of sbGLuc at the same pH, indicating increased pH resistance (bottom).

**Supplementary Figure S6**

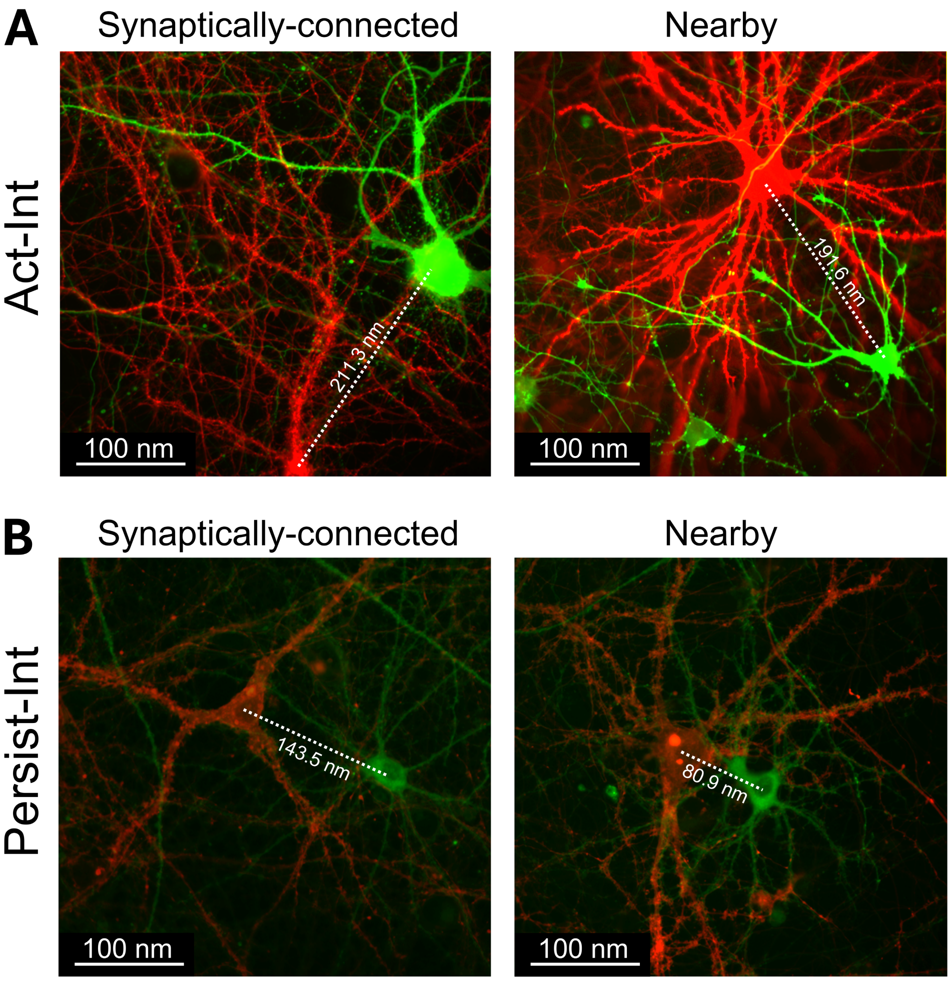

**Fig. S6. Int synaptic transmission depends on synaptic connection**. Representative images from recordings of the two modes of Interluminescence (Act-Int and Persist-Int) with pre- and postsynaptic neurons synaptically connected or non-connected (nearby). **A** Presynaptic neurons express hPOMC-sbGLuc-P2A-ChrimsonR-dTomato (red fluorescence); postsynaptic neurons express ChR2(C128S)-EYFP (green fluorescence). **B** Presynaptic neurons co-express GeNL_SS-CD4-Nrxn3b and ChrimsonR-dTomato (red fluorescence); postsynaptic neurons express ChR2(C128S)-EYFP (green fluorescence). **A,B** In all examples, opsin-expressing (green) neurons were patched. Activity was recorded after a brief red LED pulse to activate presynaptic ChrimsonR for pre- to postsynaptic endogenous neurotransmitter communication, thus determining whether the patched neuron was synaptically connected to the luciferase-expressing presynaptic neuron. This was followed by luciferin application to test for Int synaptic transmission. Changes in membrane potential (see main Figure 7E,K) were only seen in neurons confirmed to be synaptically connected. Postsynaptic neurons identified as not synaptically connected (nearby) did not show a response to luciferin application, even though nearby neurons may be closer physically in space than those synaptically connected (see indicated distances between the center of pre- and postsynaptic neurons).

#### Peptide Sequences

##### Persist-Int

>GLuc signal sequence GeNL_SS (mNGd10c-GF-d5N_SSLuc) 15aa linker ICAM Nrxn3b P2A dTomato
MGVKVLFALICIAVAEAVSKGEEDNMASLPATHELHIFGSINGVDFDMVGQGTGNPNDGYEELNLKSTKGDLQFSPW
ILVPHIGYGFHQYLPYPDGMSPFQAAMVDGSGYQVHRTMQFEDGASLTVNYRYTYEGSHIKGEAQVKGTGFPADGPV
MTNSLTAADWCRSKKTYPNDKTIISTFKWSYTTGNGKRYRSTARTTYTFAKPMAANYLKNQPMYVFRKTELKHSKTE
LNFKEWQKAFTGFEDFVGDWEQIAAYNLDQVLEQGGVSSVLQTLAVSVTPIQRIVRSGENGLKIDIHVIIPYEGLSA
DQMAHIEEVFKVVYPVDDHHFKVIMEYGTLVIDGVTPNMLNYFGRPYEGIAVFDGNKITVTGTLWNGNKIIDERLIT
PEGSMLFRVTINGVTGYRLLKKISNLEGDPLVQCGGIAGSATQTSVSPSKVILPRGGSVLVTCSTSCDQPKLLGIET
PLPKKELLLPGNNRKVYELSNVQEDSQPMCYSNCPDGQSTAKTFLTVYWTPERVELAPLPSWQPVGKNLTLRCQVEG
GAPRANLTVVLLRGEKELKREPAVGEPAEVTTTVLVRRDHHGANFSCRTELDLRPQGLELFENTSAPYQLQTFVLPA
TPPQLVSPRVLEVDTQGTVVCSLDGLFPVSEAQVHLALGDQRLNPTVTYGNDSFSAKASVSVTAEDEGTQRLTCAVI
LGNQSQETLQTVTIYSFPAPNVILTKPEVSEGTEVTVKCEAHPRAKVTLNGVPAQPLGPRAQLLLKATPEDNGRSFS
CSATLEVAGQLIHKNQTRELRVLYGPRLDERDCPGNWTWPENSQQTPMCQAWGNPLPELKCLKDGTFPLPIGESVTV
TRDLEGTYLCRARSTQGEVTRKVTVNVLSPRYESSNVASSSSTSSSPGSHSQHEHHFHGSKHHSVPISIYRSPVSLR
GGHAGATYIFGKSGGLILYTWPANDRPSTRSDRLAVGFSTTVKDGILVRIDSAPGLGAFLQLHIEQGKIGVVFNIGT
VDISIKEERTPVNDGKYHVVRFTRNGGNATLQVDNWPVNEHYPTGRQLTIFNTQAQIAIGGKDKGRLFQGQLSGLYY
DGLKVLNMAAENNPNIKINGSVRLVGEVPSILGTTQTTSMPPEMSTTVMETTTTMATTTTRKNRSTASIQPTSDDLV
SSAECSSDDEDFVECEPSTAYPYDVPDYAYPYDVPDYANPTEPGIRRVPGASEVIRESSSTTGMVVGIVAAAALCIL
ILLYAMYKYRNRDEGSYQVDETRNYISNSAQSNGTLMKEKQQSSKSGHKKQKNKDREYYVGSGATNFSLLKQAGDVE
ENPGPAAAPGPMVSKGEEVIKEFMRFKVRMEGSMNGHEFEIEGEGEGRPYEGTQTAKLKVTKGGPLPFAWDILSPQF
MYGSKAYVKHPADIPDYKKLSFPEGFKWERVMNFEDGGLVTVTQDSSLQDGTLIYKVKMRGTNFPPDGPVMQKKTMG
WEASTERLYPRDGVLKGEIHQALKLKDGGHYLVEFKTIYMAKKPVQLPGYYYVDTKLDITSHNEDYTIVEQYERSEG
RHHLFLYGMDELYK*

>GLuc signal sequence GeNL_SS (mNGd10c-GF-d5N_SSLuc) 15aa linker CD4 Nrxn3b P2A dTomato
MGVKVLFALICIAVAEAVSKGEEDNMASLPATHELHIFGSINGVDFDMVGQGTGNPNDGYEELNLKSTKGDLQFSPW
ILVPHIGYGFHQYLPYPDGMSPFQAAMVDGSGYQVHRTMQFEDGASLTVNYRYTYEGSHIKGEAQVKGTGFPADGPV
MTNSLTAADWCRSKKTYPNDKTIISTFKWSYTTGNGKRYRSTARTTYTFAKPMAANYLKNQPMYVFRKTELKHSKTE
LNFKEWQKAFTGFEDFVGDWEQIAAYNLDQVLEQGGVSSVLQTLAVSVTPIQRIVRSGENGLKIDIHVIIPYEGLSA
DQMAHIEEVFKVVYPVDDHHFKVIMEYGTLVIDGVTPNMLNYFGRPYEGIAVFDGNKITVTGTLWNGNKIIDERLIT
PEGSMLFRVTINGVTGYRLLKKISNLEGDPLVQCGGIAGSATVYKKEGEQVEFSFPLAFTVEKLTGSGELWWQAERA
SSSKSWITFDLKNKEVSVKRVTQDPKLQMGKKLPLHLTLPQALPQYAGSGNLTLALEAKTGKLHQEVNLVVMRATQL
QKNLTCEVWGPTSPKLMLSLKLENKEAKVSKREKAVWVLNPEAGMWQCLLSDSGQVLLESNIKVLPTWSTPVQPMAL
IVLGGVAGLLLFIGLGIFFCSSNVASSSSTSSSPGSHSQHEHHFHGSKHHSVPISIYRSPVSLRGGHAGATYIFGKS
GGLILYTWPANDRPSTRSDRLAVGFSTTVKDGILVRIDSAPGLGAFLQLHIEQGKIGVVFNIGTVDISIKEERTPVN
DGKYHVVRFTRNGGNATLQVDNWPVNEHYPTGRQLTIFNTQAQIAIGGKDKGRLFQGQLSGLYYDGLKVLNMAAENN
PNIKINGSVRLVGEVPSILGTTQTTSMPPEMSTTVMETTTTMATTTTRKNRSTASIQPTSDDLVSSAECSSDDEDFV
ECEPSTAYPYDVPDYAYPYDVPDYANPTEPGIRRVPGASEVIRESSSTTGMVVGIVAAAALCILILLYAMYKYRNRD
EGSYQVDETRNYISNSAQSNGTLMKEKQQSSKSGHKKQKNKDREYYVGSGATNFSLLKQAGDVEENPGPAAAPGPMV
SKGEEVIKEFMRFKVRMEGSMNGHEFEIEGEGEGRPYEGTQTAKLKVTKGGPLPFAWDILSPQFMYGSKAYVKHPAD
IPDYKKLSFPEGFKWERVMNFEDGGLVTVTQDSSLQDGTLIYKVKMRGTNFPPDGPVMQKKTMGWEASTERLYPRDG
VLKGEIHQALKLKDGGHYLVEFKTIYMAKKPVQLPGYYYVDTKLDITSHNEDYTIVEQYERSEGRHHLFLYGMDELY
K*

>GLuc signal sequence SSLuc 15aa linker B7 EYFP
MGVKVLFALICIAVAEAVFTLEDFVGDWEQIAAYNLDQVLEQGGVSSVLQTLAVSVTPIQRIVRSGENGLKIDIHVI
IPYEGLSADQMAHIEEVFKVVYPVDDHHFKVIMEYGTLVIDGVTPNMLNYFGRPYEGIAVFDGNKITVTGTLWNGNK
IIDERLITPEGSMLFRVTINGVTGYRLLKKISNGDPLVQCGGIAGSATPPEDPPDSKNTLVLFGAGFGAVITVVVIV
VIIKCFCKHRSCFRRNEASRETNNSLTFGPEEALAEQTVFLAAAVSKGEELFTGVVPILVELDGDVNGHKFSVSGEG
EGDATYGKLTLKFICTTGKLPVPWPTLVTTFGYGLQCFARYPDHMKQHDFFKSAMPEGYVQERTIFFKDDGNYKTRA

EVKFEGDTLVNRIELKGIDFKEDGNILGHKLEYNYNSHNVYIMADKQKNGIKVNFKIRHNIEDGSVQLADHYQQNTP
IGDGPVLLPDNHYLSYQSALSKDPNEKRDHMVLLEFVTAAGITLGMDELYK*

>GLuc signal sequence GeNL_eKL9h 15aa linker ICAM Nrxn3b P2A dTomato
MGVKVLFALICIAVAEAVSKGEEDNMASLPATHELHIFGSINGVDFDMVGQGTGNPNDGYEELNLKSTKGDLQFSPW
ILVPHIGYGFHQYLPYPDGMSPFQAAMVDGSGYQVHRTMQFEDGASLTVNYRYTYEGSHIKGEAQVKGTGFPADGPV
MTNSLTAADWCRSKKTYPNDKTIISTFKWSYTTGNGKRYRSTARTTYTFAKPMAANYLKNQPMYVFRKTELKHSKTE
LNFKEWQKAFTGFFEDFVGDWEQTAAYNLDQVLEQGGVSSVLQTLAVSVTPIQRIVRSGENGLKIDIHVIIPYEGLS
ADQMAHIEEVFKVVYPVDDHHFKVIMEYGTLVIDGVTPNMLNYFGRPYEGIAVFDGKKITVTGTLWNGNKIIDERLI
TPDGSMLFRVTINGVSGWRLFEKISNGDPLVQCGGIAGSATQTSVSPSKVILPRGGSVLVTCSTSCDQPKLLGIETP
LPKKELLLPGNNRKVYELSNVQEDSQPMCYSNCPDGQSTAKTFLTVYWTPERVELAPLPSWQPVGKNLTLRCQVEGG
APRANLTVVLLRGEKELKREPAVGEPAEVTTTVLVRRDHHGANFSCRTELDLRPQGLELFENTSAPYQLQTFVLPAT
PPQLVSPRVLEVDTQGTVVCSLDGLFPVSEAQVHLALGDQRLNPTVTYGNDSFSAKASVSVTAEDEGTQRLTCAVIL
GNQSQETLQTVTIYSFPAPNVILTKPEVSEGTEVTVKCEAHPRAKVTLNGVPAQPLGPRAQLLLKATPEDNGRSFSC
SATLEVAGQLIHKNQTRELRVLYGPRLDERDCPGNWTWPENSQQTPMCQAWGNPLPELKCLKDGTFPLPIGESVTVT
RDLEGTYLCRARSTQGEVTRKVTVNVLSPRYESSNVASSSSTSSSPGSHSQHEHHFHGSKHHSVPISIYRSPVSLRG
GHAGATYIFGKSGGLILYTWPANDRPSTRSDRLAVGFSTTVKDGILVRIDSAPGLGAFLQLHIEQGKIGVVFNIGTV
DISIKEERTPVNDGKYHVVRFTRNGGNATLQVDNWPVNEHYPTGRQLTIFNTQAQIAIGGKDKGRLFQGQLSGLYYD
GLKVLNMAAENNPNIKINGSVRLVGEVPSILGTTQTTSMPPEMSTTVMETTTTMATTTTRKNRSTASIQPTSDDLVS
SAECSSDDEDFVECEPSTAYPYDVPDYAYPYDVPDYANPTEPGIRRVPGASEVIRESSSTTGMVVGIVAAAALCILI
LLYAMYKYRNRDEGSYQVDETRNYISNSAQSNGTLMKEKQQSSKSGHKKQKNKDREYYVGSGATNFSLLKQAGDVEE
NPGPAAAPGPMVSKGEEVIKEFMRFKVRMEGSMNGHEFEIEGEGEGRPYEGTQTAKLKVTKGGPLPFAWDILSPQFM
YGSKAYVKHPADIPDYKKLSFPEGFKWERVMNFEDGGLVTVTQDSSLQDGTLIYKVKMRGTNFPPDGPVMQKKTMGW
EASTERLYPRDGVLKGEIHQALKLKDGGHYLVEFKTIYMAKKPVQLPGYYYVDTKLDITSHNEDYTIVEQYERSEGR
HHLFLYGMDELYK*

>GLuc signal sequence NanoLuc 15aa linker ICAM Nrxn3b P2A dTomato
MGVKVLFALICIAVAEAVFTLEDFVGDWRQTAGYNLDQVLEQGGVSSLFQNLGVSVTPIQRIVLSGENGLKIDIHVI
IPYEGLSGDQMGQIEKIFKVVYPVDDHHFKVILHYGTLVIDGVTPNMIDYFGRPYEGIAVFDGKKITVTGTLWNGNK
IIDERLINPDGSLLFRVTINGVTGWRLCERILAGDPLVQCGGIAGSATQTSVSPSKVILPRGGSVLVTCSTSCDQPK
LLGIETPLPKKELLLPGNNRKVYELSNVQEDSQPMCYSNCPDGQSTAKTFLTVYWTPERVELAPLPSWQPVGKNLTL
RCQVEGGAPRANLTVVLLRGEKELKREPAVGEPAEVTTTVLVRRDHHGANFSCRTELDLRPQGLELFENTSAPYQLQ
TFVLPATPPQLVSPRVLEVDTQGTVVCSLDGLFPVSEAQVHLALGDQRLNPTVTYGNDSFSAKASVSVTAEDEGTQR
LTCAVILGNQSQETLQTVTIYSFPAPNVILTKPEVSEGTEVTVKCEAHPRAKVTLNGVPAQPLGPRAQLLLKATPED
NGRSFSCSATLEVAGQLIHKNQTRELRVLYGPRLDERDCPGNWTWPENSQQTPMCQAWGNPLPELKCLKDGTFPLPI
GESVTVTRDLEGTYLCRARSTQGEVTRKVTVNVLSPRYESSNVASSSSTSSSPGSHSQHEHHFHGSKHHSVPISIYR
SPVSLRGGHAGATYIFGKSGGLILYTWPANDRPSTRSDRLAVGFSTTVKDGILVRIDSAPGLGAFLQLHIEQGKIGV
VFNIGTVDISIKEERTPVNDGKYHVVRFTRNGGNATLQVDNWPVNEHYPTGRQLTIFNTQAQIAIGGKDKGRLFQGQ
LSGLYYDGLKVLNMAAENNPNIKINGSVRLVGEVPSILGTTQTTSMPPEMSTTVMETTTTMATTTTRKNRSTASIQP
TSDDLVSSAECSSDDEDFVECEPSTAYPYDVPDYAYPYDVPDYANPTEPGIRRVPGASEVIRESSSTTGMVVGIVAA
AALCILILLYAMYKYRNRDEGSYQVDETRNYISNSAQSNGTLMKEKQQSSKSGHKKQKNKDREYYV*

**Act-Int**

>hPOMC(1-26) sbGLuc P2A ChrimsonR dTomato
MPRSCCSRSGALLLALLLQASMEVRGKPTENNEDFNIVAVASNFATTDLDADRGKLPGKKLPLEVLKELEANARKAG
CTRGCLICLSHIKCTPKMKKFIPGRCHTYEGDKESAQGGIGEAIVDIPEIPGFKDLEPLEQFIAQVDLCVDCTTGCL
KGLANVQCSDLLKKWLPQRCATFASKIQGQVDKIKGAGGDLEGSGATNFSLLKQAGDVEENPGPAAAATMAELISSA
TRSLFAAGGINPWPNPYHHEDMGCGGMTPTGECFSTEWWCDPSYGLSDAGYGYCFVEATGGYLVVGVEKKQAWLHSR
GTPGEKIGAQVCQWIAFSIAIALLTFYGFSAWKATCGWEEVYVCCVEVLFVTLEIFKEFSSPATVYLSTGNHAYCLR
YFEWLLSCPVILIRLSNLSGLKNDYSKRTMGLIVSCVGMIVFGMAAGLATDWLKWLLYIVSCIYGGYMYFQAAKCYV
EANHSVPKGHCRMVVKLMAYAYFASWGSYPILWAVGPEGLLKLSPYANSIGHSICDIIAKEFWTFLAHHLRIKIHEH
ILIHGDIRKTTKMEIGGEEVEVEEFVEEEDEDTVEFMVSKGEEVIKEFMRFKVRMEGSMNGHEFEIEGEGEGRPYEG
TQTAKLKVTKGGPLPFAWDILSPQFMYGSKAYVKHPADIPDYKKLSFPEGFKWERVMNFEDGGLVTVTQDSSLQDGT
LIYKVKMRGTNFPPDGPVMQKKTMGWEASTERLYPRDGVLKGEIHQALKLKDGGHYLVEFKTIYMAKKPVQLPGYYY
VDTKLDITSHNEDYTIVEQYERSEGRHHLFLYGMDELYK*

>Synaptobrevin (truncated) 15aa linker sbGLuc P2A ChrimsonR dTomato
MASQFETSAAKLKRKYWWKNLKMMIILGVICAIILIIIIVYFSTGDPLVQCGGIAGSATKPTENNEDFNIVAVASNF
ATTDLDADRGKLPGKKLPLEVLKELEANARKAGCTRGCLICLSHIKCTPKMKKFIPGRCHTYEGDKESAQGGIGEAI
VDIPEIPGFKDLEPLEQFIAQVDLCVDCTTGCLKGLANVQCSDLLKKWLPQRCATFASKIQGQVDKIKGAGGDLEGS
GATNFSLLKQAGDVEENPGPAAAATMAELISSATRSLFAAGGINPWPNPYHHEDMGCGGMTPTGECFSTEWWCDPSY
GLSDAGYGYCFVEATGGYLVVGVEKKQAWLHSRGTPGEKIGAQVCQWIAFSIAIALLTFYGFSAWKATCGWEEVYVC
CVEVLFVTLEIFKEFSSPATVYLSTGNHAYCLRYFEWLLSCPVILIRLSNLSGLKNDYSKRTMGLIVSCVGMIVFGM
AAGLATDWLKWLLYIVSCIYGGYMYFQAAKCYVEANHSVPKGHCRMVVKLMAYAYFASWGSYPILWAVGPEGLLKLS
PYANSIGHSICDIIAKEFWTFLAHHLRIKIHEHILIHGDIRKTTKMEIGGEEVEVEEFVEEEDEDTVEFMVSKGEEV
IKEFMRFKVRMEGSMNGHEFEIEGEGEGRPYEGTQTAKLKVTKGGPLPFAWDILSPQFMYGSKAYVKHPADIPDYKK
LSFPEGFKWERVMNFEDGGLVTVTQDSSLQDGTLIYKVKMRGTNFPPDGPVMQKKTMGWEASTERLYPRDGVLKGEI
HQALKLKDGGHYLVEFKTIYMAKKPVQLPGYYYVDTKLDITSHNEDYTIVEQYERSEGRHHLFLYGMDELYK*

>VGAT sbGLuc P2A ChrimsonR dTomato
MATLLRSKLSNVATSVSNKSQAKMSGMFARMGFQAATDEEAVGFAHCDDLDFEHRQGLQMDILKAEGEPCGDEGAEA
PVEGDIHYQRGSGAPLPPSGSKDQVGGGGEFGGHDKPKITAWEAGWNVTNAIQGMFVLGLPYAILHGGYLGLFLIIF
AAVVCCYTGKILIACLYEENEDGEVVRVRDSYVAIANACCAPRFPTLGGRVVNVAQIIELVMTCILYVVVSGNLMYN
SFPGLPVSQKSWSIIATAVLLPCAFLKNLKAVSKFSLLCTLAHFVINILVIAYCLSRARDWAWEKVKFYIDVKKFPI
SIGIIVFSYTSQIFLPSLEGNMQQPSEFHCMMNWTHIAACVLKGLFALVAYLTWADETKEVITDNLPGSIRAVVNIF
LVAKALLSYPLPFFAAVEVLEKSLFQEGSRAFFPACYSGDGRLKSWGLTLRCALVVFTLLMAIYVPHFALLMGLTGS
LTGAGLCFLLPSLFHLRLLWRKLLWHQVFFDVAIFVIGGICSVSGFVHSLEGLIEAYRTNAEDKPTENNEDFNIVAV
ASNFATTDLDADRGKLPGKKLPLEVLKELEANARKAGCTRGCLICLSHIKCTPKMKKFIPGRCHTYEGDKESAQGGI
GEAIVDIPEIPGFKDLEPLEQFIAQVDLCVDCTTGCLKGLANVQCSDLLKKWLPQRCATFASKIQGQVDKIKGAGGD
LEGSGATNFSLLKQAGDVEENPGPAAAATMAELISSATRSLFAAGGINPWPNPYHHEDMGCGGMTPTGECFSTEWWC
DPSYGLSDAGYGYCFVEATGGYLVVGVEKKQAWLHSRGTPGEKIGAQVCQWIAFSIAIALLTFYGFSAWKATCGWEE
VYVCCVEVLFVTLEIFKEFSSPATVYLSTGNHAYCLRYFEWLLSCPVILIRLSNLSGLKNDYSKRTMGLIVSCVGMI
VFGMAAGLATDWLKWLLYIVSCIYGGYMYFQAAKCYVEANHSVPKGHCRMVVKLMAYAYFASWGSYPILWAVGPEGL
LKLSPYANSIGHSICDIIAKEFWTFLAHHLRIKIHEHILIHGDIRKTTKMEIGGEEVEVEEFVEEEDEDTVEFMVSK
GEEVIKEFMRFKVRMEGSMNGHEFEIEGEGEGRPYEGTQTAKLKVTKGGPLPFAWDILSPQFMYGSKAYVKHPADIP
DYKKLSFPEGFKWERVMNFEDGGLVTVTQDSSLQDGTLIYKVKMRGTNFPPDGPVMQKKTMGWEASTERLYPRDGVL
KGEIHQALKLKDGGHYLVEFKTIYMAKKPVQLPGYYYVDTKLDITSHNEDYTIVEQYERSEGRHHLFLYGMDELYK*

>hPOMC(1-26) mCherry22.0_RLuc8.6(W535/F156)
MPRSCCSRSGALLLALLLQASMEVRGVSKGEEDNMAIIKEYMRFKVHMEGSVNGHEFEIEGEGEGRPFEGTQTAKLK
VTKGGPLPFAWHILPPQFQYGSKAYVKHPADIPDYFKLSFPEGFTWEREMNFEDGGVVTVTQDSSLQDGEFIYKVKL
RGTNFPSDGPVMQKKTMGNTASTERMYPEDGALKGETKWRLKLKDGGHYEAEVKTTYKAKKPVQLPGAYNVDRKLDI
TYHNEDYTIVEQYERAEARHIDKVYDPEQRKRMITGPQWWARCKQMNVLDSFINYYDSEKHAENAVIFLHGNATSSY
LWRHVVPHIEPVARCIIPDLIGMGKSGKSGNGSYRLLDHYKYLTAWFELLNLPKKIIFVGHDWGSALAFHYAYEHQD
RIKAIVHMESVVDVIESWMGFPDIEEELALIKSEEGEKMVLENNFFVETLLPSKIMRKLEPEEFAAYLEPFKEKGEV
RRPTLSWPREIPLVKGGKPDVVQIVRNYNAYLRASDDLPKLFIESDPGFFSNAIVEGAKKFPNTEFVKVKGLHFLQE
DAPDEMGKYIKSFVERVLKNEQ*

>hPOMC(1-26) GeNL_SS (mNGd10c-GF-d5N_SSLuc) P2A ChrimsonR dTomato
MPRSCCSRSGALLLALLLQASMEVRGVSKGEEDNMASLPATHELHIFGSINGVDFDMVGQGTGNPNDGYEELNLKST
KGDLQFSPWILVPHIGYGFHQYLPYPDGMSPFQAAMVDGSGYQVHRTMQFEDGASLTVNYRYTYEGSHIKGEAQVKG
TGFPADGPVMTNSLTAADWCRSKKTYPNDKTIISTFKWSYTTGNGKRYRSTARTTYTFAKPMAANYLKNQPMYVFRK
TELKHSKTELNFKEWQKAFTGFEDFVGDWEQIAAYNLDQVLEQGGVSSVLQTLAVSVTPIQRIVRSGENGLKIDIHV
IIPYEGLSADQMAHIEEVFKVVYPVDDHHFKVIMEYGTLVIDGVTPNMLNYFGRPYEGIAVFDGNKITVTGTLWNGN
KIIDERLITPEGSMLFRVTINGVTGYRLLKKISNLEGSGATNFSLLKQAGDVEENPGPAAAATMAELISSATRSLFA
AGGINPWPNPYHHEDMGCGGMTPTGECFSTEWWCDPSYGLSDAGYGYCFVEATGGYLVVGVEKKQAWLHSRGTPGEK
IGAQVCQWIAFSIAIALLTFYGFSAWKATCGWEEVYVCCVEVLFVTLEIFKEFSSPATVYLSTGNHAYCLRYFEWLL
SCPVILIRLSNLSGLKNDYSKRTMGLIVSCVGMIVFGMAAGLATDWLKWLLYIVSCIYGGYMYFQAAKCYVEANHSV
PKGHCRMVVKLMAYAYFASWGSYPILWAVGPEGLLKLSPYANSIGHSICDIIAKEFWTFLAHHLRIKIHEHILIHGD
IRKTTKMEIGGEEVEVEEFVEEEDEDTVEFMVSKGEEVIKEFMRFKVRMEGSMNGHEFEIEGEGEGRPYEGTQTAKL
KVTKGGPLPFAWDILSPQFMYGSKAYVKHPADIPDYKKLSFPEGFKWERVMNFEDGGLVTVTQDSSLQDGTLIYKVK
MRGTNFPPDGPVMQKKTMGWEASTERLYPRDGVLKGEIHQALKLKDGGHYLVEFKTIYMAKKPVQLPGYYYVDTKLD
ITSHNEDYTIVEQYERSEGRHHLFLYGMDELYK*

>hPOMC(1-26) SSLuc P2A ChrimsonR dTomato
MPRSCCSRSGALLLALLLQASMEVRGVFTLEDFVGDWEQIAAYNLDQVLEQGGVSSVLQTLAVSVTPIQRIVRSGEN
GLKIDIHVIIPYEGLSADQMAHIEEVFKVVYPVDDHHFKVIMEYGTLVIDGVTPNMLNYFGRPYEGIAVFDGNKITV
TGTLWNGNKIIDERLITPEGSMLFRVTINGVTGYRLLKKISNLEGSGATNFSLLKQAGDVEENPGPAAAATMAELIS
SATRSLFAAGGINPWPNPYHHEDMGCGGMTPTGECFSTEWWCDPSYGLSDAGYGYCFVEATGGYLVVGVEKKQAWLH
SRGTPGEKIGAQVCQWIAFSIAIALLTFYGFSAWKATCGWEEVYVCCVEVLFVTLEIFKEFSSPATVYLSTGNHAYC
LRYFEWLLSCPVILIRLSNLSGLKNDYSKRTMGLIVSCVGMIVFGMAAGLATDWLKWLLYIVSCIYGGYMYFQAAKC
YVEANHSVPKGHCRMVVKLMAYAYFASWGSYPILWAVGPEGLLKLSPYANSIGHSICDIIAKEFWTFLAHHLRIKIH
EHILIHGDIRKTTKMEIGGEEVEVEEFVEEEDEDTVEFMVSKGEEVIKEFMRFKVRMEGSMNGHEFEIEGEGEGRPY
EGTQTAKLKVTKGGPLPFAWDILSPQFMYGSKAYVKHPADIPDYKKLSFPEGFKWERVMNFEDGGLVTVTQDSSLQD
GTLIYKVKMRGTNFPPDGPVMQKKTMGWEASTERLYPRDGVLKGEIHQALKLKDGGHYLVEFKTIYMAKKPVQLPGY
YYVDTKLDITSHNEDYTIVEQYERSEGRHHLFLYGMDELYK*

>hPOMC(1-26) GGS linker SSLuc_smb P2A ChrimsonR dTomato
MPRSCCSRSGALLLALLLQASMEVRGGGSGGSGGVTGYRLFEEILLEGSGATNFSLLKQAGDVEENPGPAAAATMAE
LISSATRSLFAAGGINPWPNPYHHEDMGCGGMTPTGECFSTEWWCDPSYGLSDAGYGYCFVEATGGYLVVGVEKKQA
WLHSRGTPGEKIGAQVCQWIAFSIAIALLTFYGFSAWKATCGWEEVYVCCVEVLFVTLEIFKEFSSPATVYLSTGNH
AYCLRYFEWLLSCPVILIRLSNLSGLKNDYSKRTMGLIVSCVGMIVFGMAAGLATDWLKWLLYIVSCIYGGYMYFQA
AKCYVEANHSVPKGHCRMVVKLMAYAYFASWGSYPILWAVGPEGLLKLSPYANSIGHSICDIIAKEFWTFLAHHLRI
KIHEHILIHGDIRKTTKMEIGGEEVEVEEFVEEEDEDTVEFMVSKGEEVIKEFMRFKVRMEGSMNGHEFEIEGEGEG
RPYEGTQTAKLKVTKGGPLPFAWDILSPQFMYGSKAYVKHPADIPDYKKLSFPEGFKWERVMNFEDGGLVTVTQDSS
LQDGTLIYKVKMRGTNFPPDGPVMQKKTMGWEASTERLYPRDGVLKGEIHQALKLKDGGHYLVEFKTIYMAKKPVQL
PGYYYVDTKLDITSHNEDYTIVEQYERSEGRHHLFLYGMDELYK*

>GLuc signal sequence SSLuc_lgb GGS linker 15aa linker ChRmine EYFP
MGVKVLFALICIAVAEAVFTLEDFVGDWEQIAAYNLDQVLEQGGVSSVLQTLAVSVTPIQRIVRSGENGLKIDIHVI
IPYEGLSADQMAHIEEVFKVVYPVDDHHFKVIMEYGTLVIDGVTPNMLNYFGRPYEGIAVFDGKKITVTGTLWNGNK
IIDERLITPEGSMLFRVTINSGGSGGSGGLEGDPLVQCGGIAGSATAHAPGTDQMFYVGTMDGWYLDTKLNSVAIGA
HWSCFIVLTITTFYLGYESWTSRGPSKRTSFYAGYQEEQNLALFVNFFAMLSYFGKIVADTLGHNFGDVGPFIIGFG
NYRYADYMLTCPMLVYDLLYQLRAPYRVSCSAIIFAILMSGVLAEFYAEGDPRLRNGAYAWYGFGCFWFIFAYSIVM
SIVAKQYSRLAQLAQDTGAEHSLHVLKFAVFTFSMLWILFPLVWAICPRGFGWIDDNWTEVAHCVCDIVAKSCYGFA
LARFRKTYDEELFRLLEQLGHDEDEFQKLELDMRLSSNGERLRRLSAAAKSRITSEGEYIPLDQIDINVVSKGEELF
TGVVPILVELDGDVNGHKFSVSGEGEGDATYGKLTLKFICTTGKLPVPWPTLVTTFGYGLQCFARYPDHMKQHDFFK
SAMPEGYVQERTIFFKDDGNYKTRAEVKFEGDTLVNRIELKGIDFKEDGNILGHKLEYNYNSHNVYIMADKQKNGIK
VNFKIRHNIEDGSVQLADHYQQNTPIGDGPVLLPDNHYLSYQSALSKDPNEKRDHMVLLEFVTAAGITLGMDELYKF
CYENEV*
